# Start codon disruption with CRISPR/Cas9 prevents murine Fuchs’ endothelial corneal dystrophy

**DOI:** 10.1101/2020.03.18.996728

**Authors:** Hironori Uehara, Xiaohui Zhang, Felipe Pereira, Siddharth Narendran, Susie Choi, Sai Bhuvanagiri, Jinlu Liu, Sangeetha Ravi Kumar, Austin Bohner, Lara Carroll, Bonnie Archer, Yue Zhang, Wei Liu, Guangping Gao, Jayakrishna Ambati, Albert S Jun, Balamurali K. Ambati

## Abstract

A missense mutation of collagen type VIII alpha 2 chain (COL8A2) gene leads to early onset Fuchs’ endothelial corneal dystrophy (FECD), which progressively impairs vision through loss of corneal endothelial cells. We demonstrate that CRISPR/Cas9-based postnatal gene editing achieves structural and functional rescue in a mouse model of FECD. A single intraocular injection of an adenovirus encoding both the Cas9 gene and guide RNA (Ad-Cas9-Col8a2gRNA), efficiently knocked down mutant COL8A2 expression in corneal endothelial cells, prevented endothelial cell loss, and rescued corneal endothelium pumping function in adult Col8a2 mutant mice. There were no adverse sequelae on histology or electroretinography. Col8a2 start codon disruption represents a non-surgical strategy to prevent vision loss in early-onset FECD. As this demonstrates the ability of Ad-Cas9-gRNA to restore phenotype in adult post-mitotic cells, this method may be widely applicable to adult-onset diseases, even in tissues affected with disorders of non-reproducing cells.

## Introduction

Fuchs’ endothelial corneal dystrophy (FECD), which is characterized by progressive loss of corneal endothelial cells, is the leading cause of corneal transplantation in industrialized societies^1^. Currently, the only available treatment for advanced FECD is corneal transplantation, which entails significant risks (e.g. infection, hemorrhage, rejection, glaucoma) both during surgery and the lifetime of the patient^2,3^. A missense mutation of the collagen 8A2 (*COL8A2*) gene in humans causes early onset Fuchs’ dystrophy^4,5,6^. Although other mutations within the *ZEB1/TCF8* locus and *TCF4* trinucleotide repeats are associated with Fuchs’ dystrophy^7–15^, only the *Col8a2* missense mutant mouse has successfully recapitulated its key features. Two distinct transgenic approaches in mice have helped illuminate the role of *Col8a2* in the onset of FECD. Knockout mice lacking *Col8a2* alone or combined with a homozygous *Col8a1* knockout mutation do not develop FECD^16^. Although the double knockouts exhibited corneal biomechanical weakening (without endothelial loss), *Col8a2* knockouts showed no apparent phenotype. In contrast, *Col8a2* mutant knock-in mice carrying the Q455K and L450W mutations associated with early-onset FECD in human patients,, displayed corneal endothelial excrescences known as guttae, as well as the endothelial cell loss that are hallmarks of human FECD^17,18^. Taken together, these studies suggest that *COL8A2* protein is not essential to corneal function yet is causally responsible for FECD via mutant dominant gain-of-function activity. We therefore sought to test whether knock-down of mutant *COL8A2* could offer a new therapeutic strategy for early-onset FECD, establishing a precedent for treating gain-of-function genetic disorders in post-mitotic cells by tissue-specific ablation of the missense gene, targeting the start codon with CRISPR/Cas9.

## Results

### Strategy of mouse Col8a2 gene knock down by CRISPR/Cas9

To disrupt *Col8a2* gene expression, we designed a guide RNA (gRNA) targeting the start codon of the *Col8a2* gene (MsCol8a2gRNA) by non-homologous end-joint repair through CRISPR/Cas9^19,20^ (**Figure 1a**). The strategy of targeting the start codon is sufficient for blocking gene expression at the translational level. The appeal of this strategy, as opposed to correcting the mutation through homologous recombination (HR), is that poor efficiency of CRISPR-based HR would result in a majority of sequence changes comprised of insertion/deletions (indel). Consequently, the further one targets downstream from the start codon, the greater the risk of missense mutations that result in viable mutant proteins with unknown activity. By targeting inside or near the start codon, this risk is minimized. As a backbone plasmid, we used pX330-U6-Chimeric_BB-CBh-hSpCas9^20^ which encodes spCas9 and gRNA downstream of the U6 promoter (px330-MsCol8a2gRNA1). To detect the indel, we used CviAII or Hin1II digestion of PCR products (**Figure 1b**). CviAII/Hin1II cuts 5’-CATG-3’, which digests at the *Col8A2* start codon, whereas an undigested band indicates the presence of an indel at the start codon. As expected, px330-MsCol8a2gRNA1 creates an indel in mouse NIH3T3 cells (**Figure 1b**). Furthermore, we designed MsCol8a2gRNA2 and MsCol8a2gRNA3 downstream of MsCol8a2gRNA1 (**Figure 1a**). Co-transfection of px330-MsCol8a2gRNA1 with px330-MsCol8a2gRNA2 or px330-MsCol8a2gRNA3 resulted in an extra PCR band (**Figure 1c**). The indels by px330-MsCol8A2gRNA1 were confirmed by sequencing (**Figure 1d**). Although two guide RNAs could potentially attenuate target gene expression more efficiently than a single guide RNA, we proceeded with *in vivo* experiments using only MsCol8a2gRNA1.

**Figure 1.**
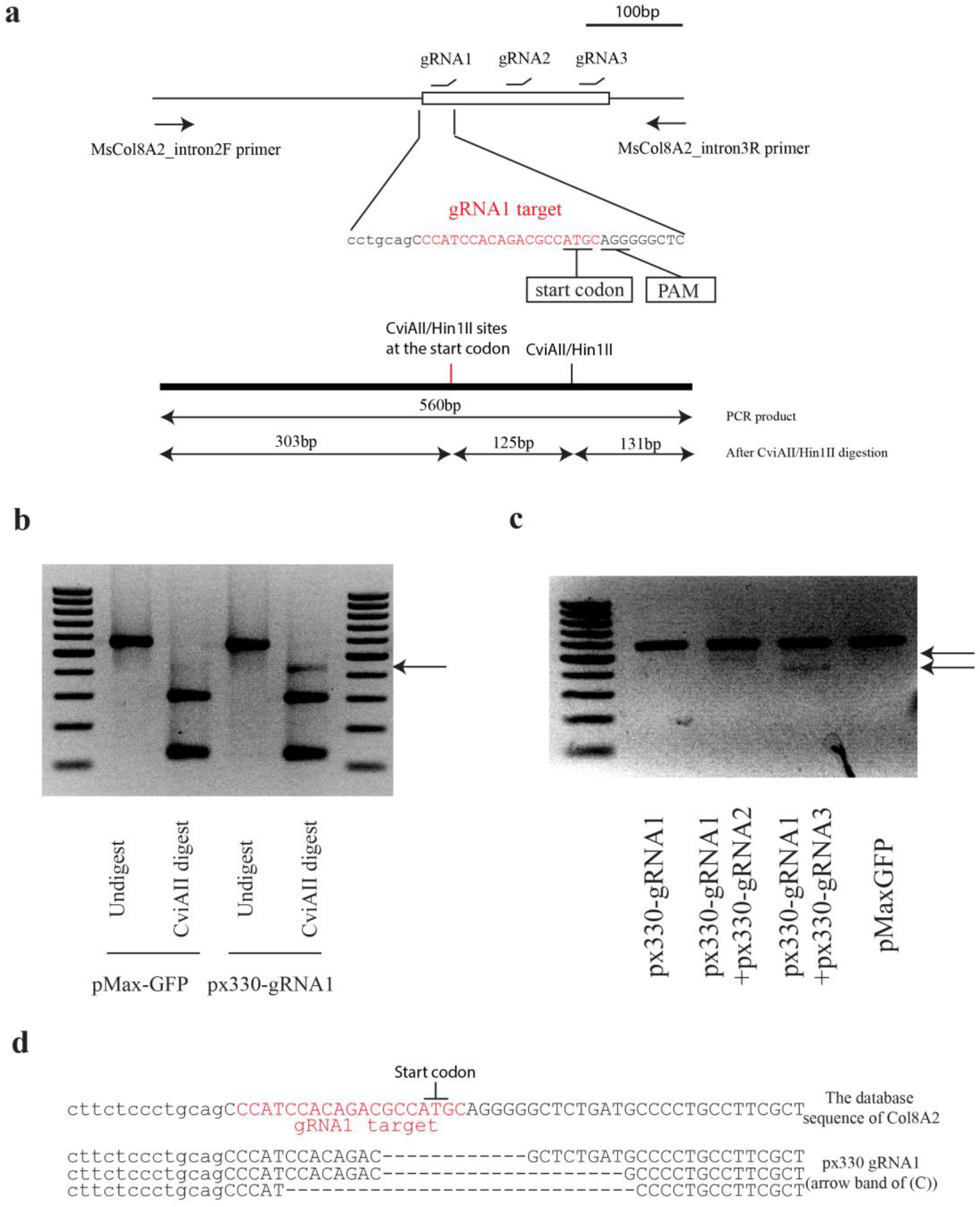
Design of *Col8a2* guide RNA and indel confirmation *in vitro*. **a,** Design of gRNAs for mouse *Col8a2* gene and schematic diagram of indel detection by restriction enzyme digestion of PCR product. gRNA1, which is used for Ad-Cas9-Col8a2gRNA, was designed to disrupt the *Col8a2* start codon. PCR primers were designed to flank the start codon and gRNA targeting sites. PCR product from the intact DNA sequence is 560bp, which is digested to 303bp, 131bp and 126bp by CviAII/Hin1II restriction enzymes. **b,** In px330-gRNA1 transfected NIH3T3 cells, the PCR product showed an extra band (~430 bp, arrow) after CviAII digestion. pMax-GFP was used as a control. **c,** Combination of two plasmids (px330-gRNA1+px330-gRNA2 and px330-gRNA1+px330-gRNA2) yields lower bands (arrow) reflecting the deletion between the targeted sites. **d,** The deletion of the start codon by px330-gRNA1 was confirmed by Sanger sequencing after cloning.

### *In vivo Col8a2* gene knock down in mouse corneal endothelium by adenovirus mediated CRISPR/Cas9

To introduce the genes (SpCas9 and gRNA) into corneal endothelium in vivo, we produced recombinant adenovirus Cas9-Col8a2gRNA (Ad-Cas9-Col8a2gRNA). There are several common viruses such as adeno-associated virus and lentivirus, but previous studies indicated only adenovirus has efficient gene transfer to corneal endothelium in vivo. In fact, we found adenovirus-GFP showed efficient GFP expression in corneal endothelium (**Figure 2a**). First, we determined the effective adenovirus dose *in vitro* for indel production at the *Col8a2* start codon (**Supplemental Figure 1a-c**). To confirm effective indel production *in vivo*, we tested various titers of Ad-Cas9-Col8a2gRNA injected into the aqueous humor of adult C57BL/6J mice. After one month, the corneal endothelium/stroma and epithelium/stroma were separated mechanically (**Supplemental Figure 2a-f**), followed by genomic DNA purification. Digestion of PCR products by CviAII/Hin1II revealed an undigested band from amplified corneal endothelium DNA (arrow in **Figure 2b**) indicating disruption of the *Col8a2* start codon, which was confirmed by sanger sequence analysis (**Figure 2c**). In contrast, corneal epithelium and stroma revealed an intact start codon after CviAII/Hin1II digestion of PCR amplified DNA.

**Figure 2.**
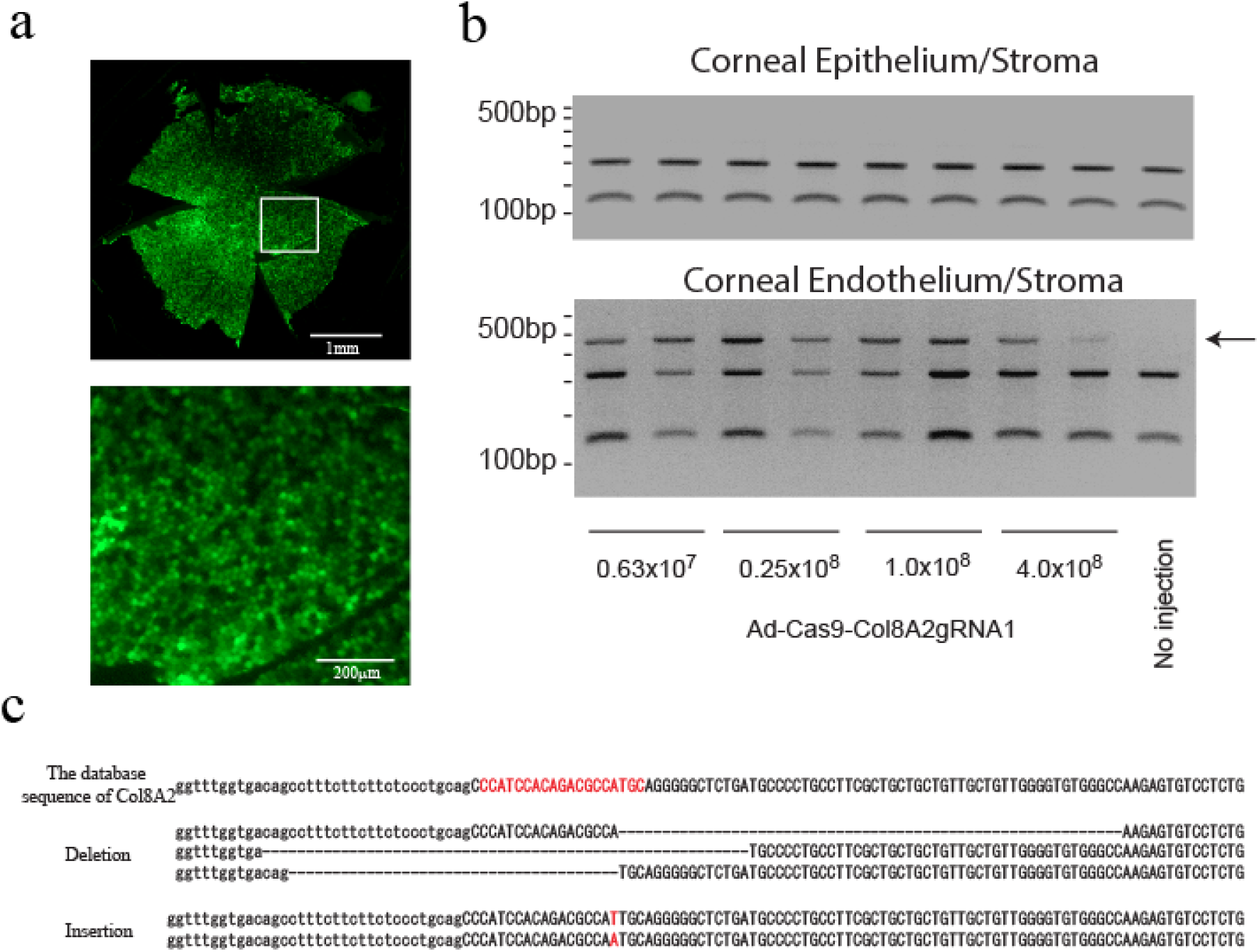
Intracameral injection of Ad-Cas9-Col8a2gRNA1 induces indel at the Col8a2 start codon in corneal endothelium. **a,** Adenovirus infection to corneal endothelium via intracameral injection was confirmed by adenovirus GFP. Top: whole mouse cornea flatmount. Bottom: the magnified image. **b,** Ad-Cas9-Col8a2gRNA1 induces an insertion/deletion (indel) at the *Col8a2* start codon in the corneal endothelium but not in the corneal epithelium/stroma. Genomic DNA of corneal endothelium/stroma and corneal epithelium/stroma was PCR amplified with primers flanking the *Col8a2* start site and digested with CviAII, which recognizes the intact *Col8a2* start codon (5’-CATG-3’). The CviAII undigested band (arrow) demonstrates the indel at the *Col8a2* start codon. **c,** Sanger sequencing of the cloned PCR product from genomic DNA purified from corneal endothelium/stroma confirm indels at the *Col8a2* start codon.

Next, to examine whether start codon disruption reduces COL8A2 protein expression in the corneal endothelium, we localized protein in sectioned corneas with an anti-COL8A2 antibody (**Figure 3** and **Supplemental Figure 3a-e**). The non-injected cornea showed COL8A2 protein expression in corneal epithelium and endothelium. As predicted, Ad-Cas9-Col8a2gRNA injected corneas exhibited reduced COL8A2 protein expression in corneal endothelium but not corneal epithelium. Thus, we successfully knocked down *in vivo* COL8A2 protein expression in adult corneal endothelium by Ad-Cas9-Col8a2gRNA.

**Figure 3.**
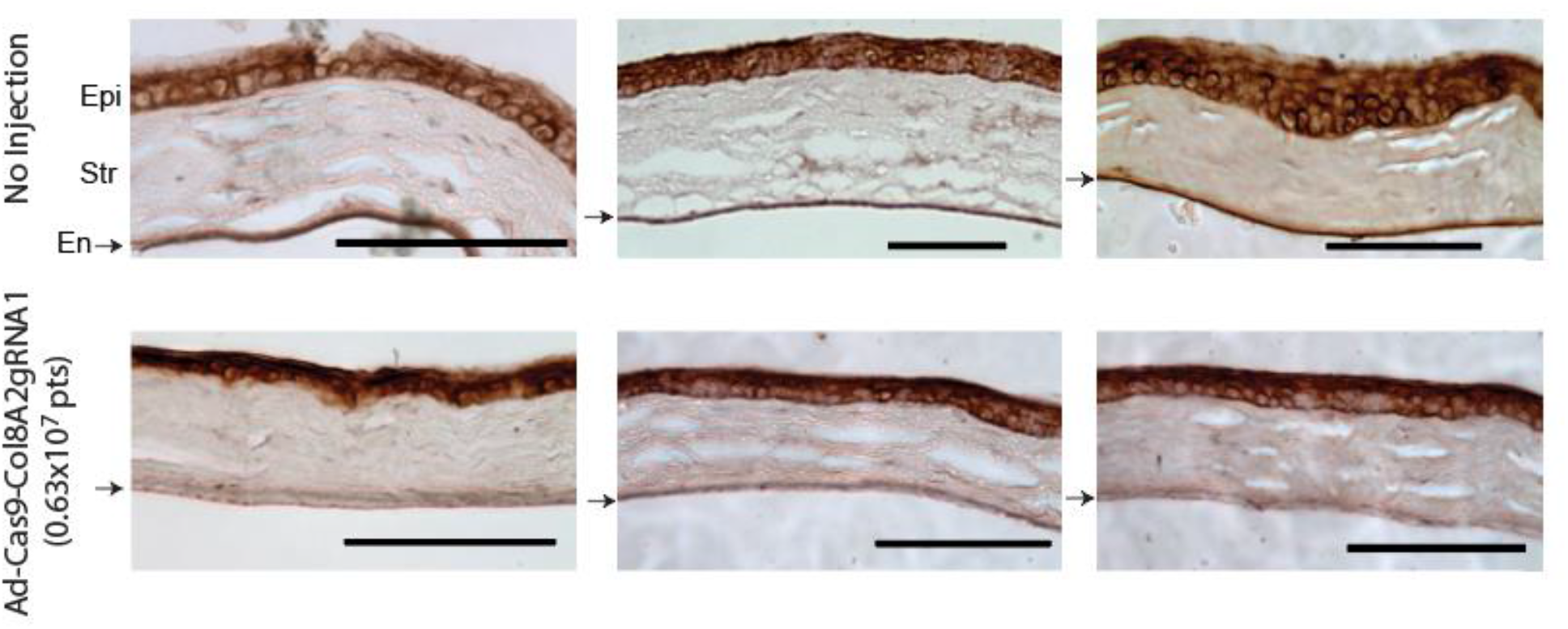
Ad-Cas9-Col8a2gRNA reduces COL8A2 expression in mouse corneal endothelium but not epithelium. COL8A2 protein immunostaining from the cornea two months after injection with DPBS (4µL, upper figures) or Ad-Cas9-Col8a2gRNA (0.63 × 10^7^vg in 4µL, lower figures). In Ad-Cas9-Col8a2gRNA injected corneas, lower COL8A2 protein expression was seen in corneal endothelium, but not in epithelium. Epi: epithelium, Str: stroma, En (arrow): endothelium. Scale bar = 100µm.

### Determination of the Safety dose of Ad-Cas9-Col8a2gRNA

As adenoviruses are known to induce inflammation and cell toxicity, we tested a range of Ad-Cas9-Col8a2gRNA titers for safety. Anterior chamber injection of the highest titer (4.0 × 10^8^vg) devastated the mouse corneal endothelium, inducing corneal opacity and edema in C57BL/6J mice (**Supplemental Figure 4**). Although corneal thickness and histopathology appeared normal at lower titers (**Supplemental Figures 5-7**), ZO-1 immunolabeling detected reduced endothelial density in corneal flat mounts after injecting 1.0 × 10^8^vg (**Supplemental Figure 8**). At 0.25 × 10^8^vg, neither tumor necrosis factor alpha (TNFα) nor interferon gamma (IFNγ) were upregulated 4 weeks after Ad-Cas9-Col8a2gRNA injection (**Supplemental Figure 9**). Moreover, we confirmed that Ad-Cas9-Col8a2gRNA did not suppress retinal function, as monitored by electroretinography (ERG), or damage retinal structure, as visualized by hematoxylin-eosin (HE) staining a (**Supplemental Figure 10 and 11a**). Finally, anterior chamber injection of Ad-Cas9-Col8a2gRNA did not induce liver or kidney damage or inflammation, as visualized by HE staining (**Supplemental Figure 11b**). Hence, subsequent experiments were performed with 0.25 × 10^8^vg of Ad-Cas9-Col8a2gRNA, which did not induce detectable toxicity.

### Efficiency of indel induction by Ad-Cas9-Col8a2gRNA *in vivo*

To determine the indel rate in mouse corneal endothelium, we performed deep sequencing of PCR products (including the target site) amplified from genomic DNA of corneal endothelium. We found that the indel rate was 23.7 ± 4.5% in mouse corneal endothelium (**Table 1**). Most insertions were 1bp insertions (19.8 ± 4.0% in total reads, **Figure 4a**), while 2bp deletions were the most frequent (1.0 ± 0.3% in total reads, **Figure 4b**). We moreover found that A or T insertion was predominant, with the proportion of A:T:G:C being 48.7 : 44.6 : 1.8 : 4.9 (**Table 2**). Adenine insertion (9.4 ± 1.9% in total reads) produced a cryptic ATG start codon (**Supplemental Figure 12**). This insertion changes G to C at the -3 position (A in ATG as +1). Since previous studies indicated G or A at the -3 position is important for translational commencement which is known as a kozak sequence^21,22^, a consequent reduction in protein expression by the disruption of kozak/ATG sequence would be predicted.

**Figure 4.**
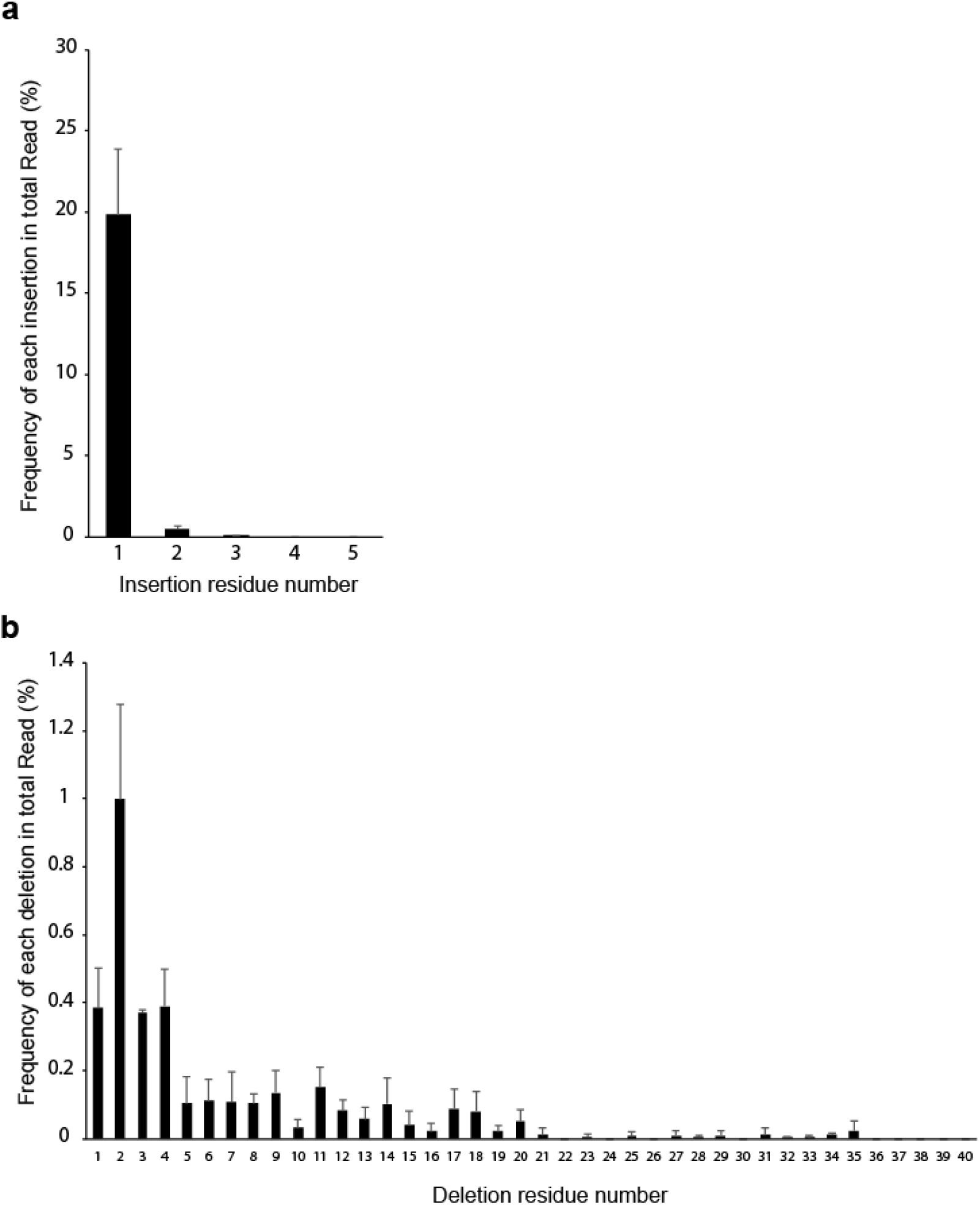
Distribution of inserted and deleted residue number. **a,** Frequency of insertion. 1bp insertion was most frequent. **b,** Frequency of deletion. 2bp deletion was most frequent. n=4. Error bar is standard deviation.

**Table 1.**
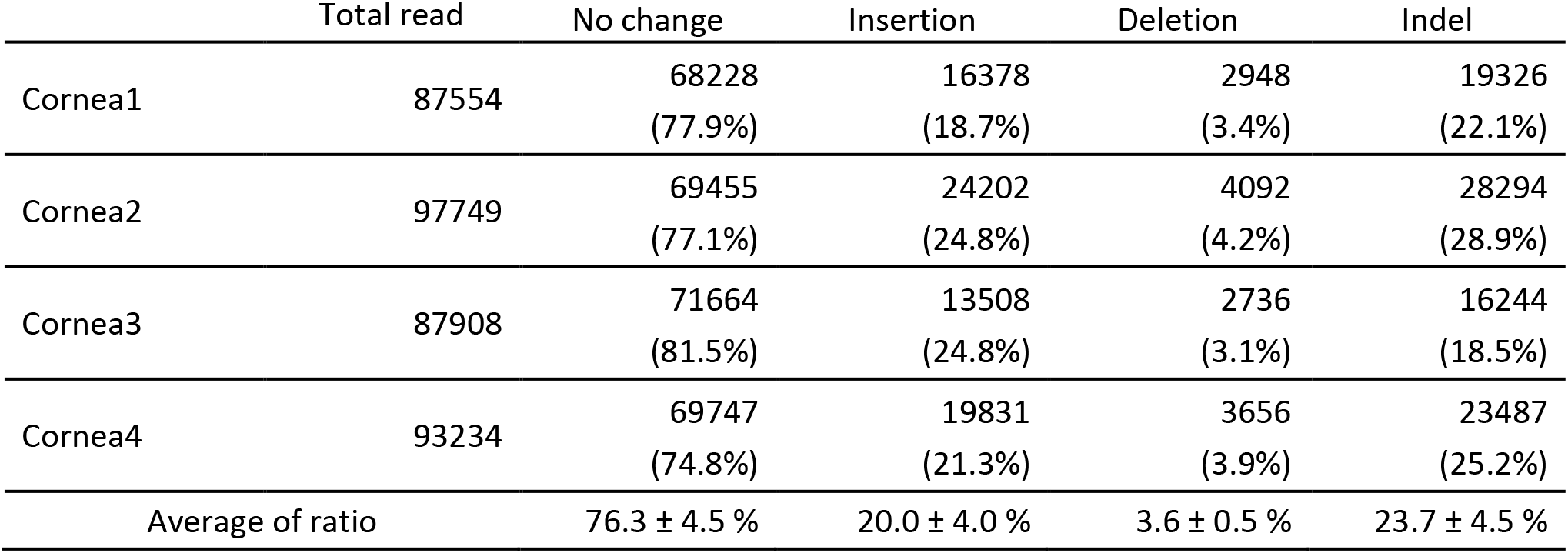
Indel rate at mouse Col8a2 target site by Ad-Cas9-Col8a2gRNA from corneal endothelium.

**Table 2.**
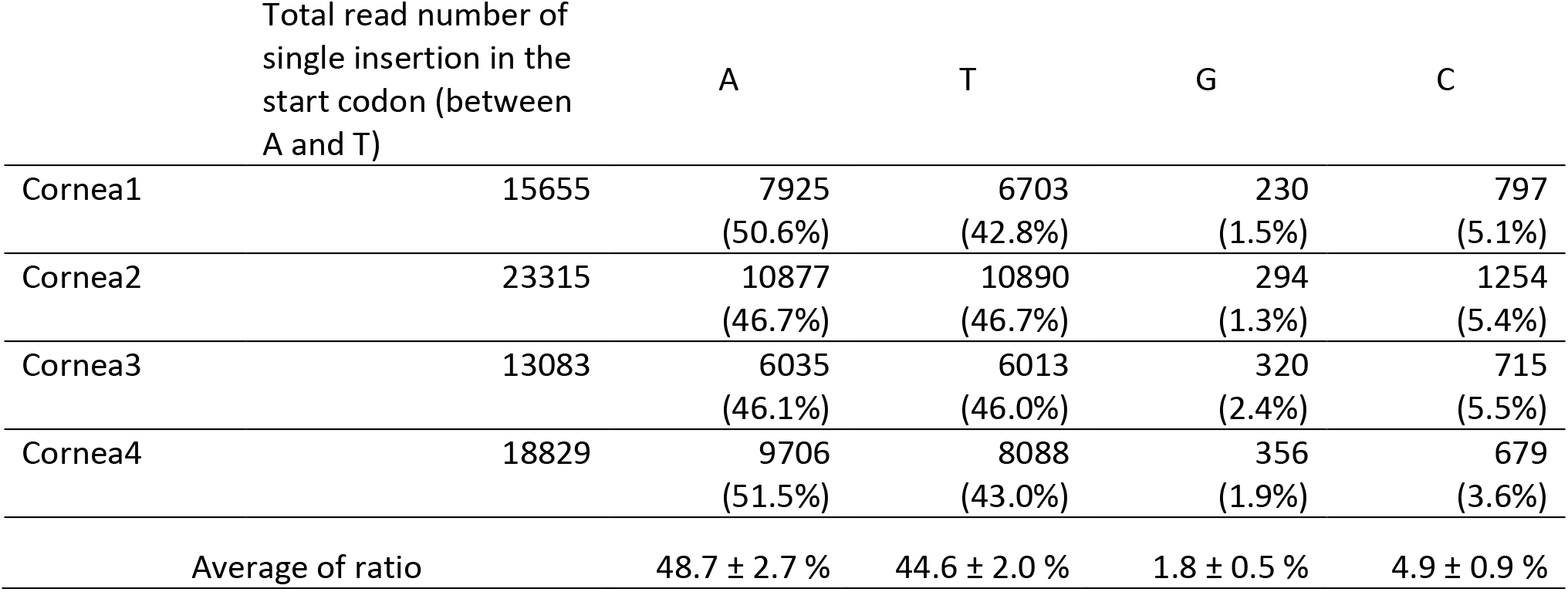
Ratio of A:T:G:C in 1 bp insertions.

The indel rate in corneal endothelium was 23.7 ± 4.5%, which was much lower than the anticipated since COL8A2 protein expression in mouse corneal endothelium was markedly decreased by anterior chamber injection of Ad-Cas9-Col8a2gRNA (**Figure 3 and Supplemental Figure 3**) and because of the high rate of adenovirus infection of the corneal endothelium (**Figure 2a**). We speculate this is due to gDNA from corneal stroma cells based on the following. The number of corneal endothelial cells is approximately 7200 cells (2300 cells/mm^2^ × 1mm × 1mm × π), and then the expected purified gDNA amount is 43 ng as a genome mass from mouse cell is 6 pg ((5.46 × 10^9^ as 2n) × 660 (average molecular weight of DNA base pair) / (6.02 × 10^−23^, Avogadròs number)). The purified gDNA from the peeled endothelium was a much higher amount than predicted (**Table 3**). We therefore hypothesized that stromal cells were contained in our samples. To confirm this, we conducted experiments in **Supplemental figure 2**. We peeled half of corneal endothelium, placed back in situ, and then proceeded to cryosection with DAPI staining. As we expected, we found many stroma cells were found with corneal endothelium. Hence, the extra gDNA is stromal-derived. Therefore, we can normalize indel rate by the proportion of endothelial cell gDNA to total isolated gDNA (**Table 3**). From this calculation, the normalized indel rate (proportion of endothelial cells with indels) is 102.5 ±16.3 %. This accords with the observed immunostaining pattern in **Figure 3**.

**Table 3.** Normalized Indel rate by the purified genomic DNA amount.

|  | Concentraion<br>(ng/uL) | gDNA amount<br>(ng, 16uL<br>elution) | Cell number<br>from gDNA<br>amount | Intact Indel rate<br>(%) | Normalized<br>Indel rate (%) |
| --- | --- | --- | --- | --- | --- |
| Cornea1 | 14.5 | 232 | 38744 | 22.1 | 118.6 |
| Cornea2 | 10.5 | 168 | 28056 | 28.9 | 112.3 |
| Cornea3 | 12 | 192 | 32064 | 18.5 | 82.1 |
| Cornea4 | 10.4 | 166.4 | 27789 | 25.2 | 97.0 |

### Ad-Cas9-Col8a2gRNA rescues corneal endothelium architecture in *Col8a2*^Q455K/Q455K^ FECD mice

Next, we examined whether Ad-Cas9-Col8a2gRNA rescued corneal endothelium in the early-onset *Col8a2*^Q455K/Q455K^ FECD mouse model^18^. At two months of age, we performed a single intraocular injection of Ad-Cas9-Col8a2gRNA into one eye. Uninjected contralateral eyes were used as controls. After the injection, the corneal endothelium was examined by *in vivo* corneal confocal microscopy (**Figure 5a**). Ad-Cas9-Col8a2gRNA injected eyes showed slower reduction of corneal endothelium than the uninjected eyes (**Figure 5b**). After 10 months (12-month-old), apparent differences between corneal endothelium of Ad-Cas9-Col8a2gRNA injected and uninjected eyes were obvious (**Figure 5c**). We found that intraocular injection of Ad-Cas9-Col8a2gRNA significantly rescued corneal endothelium in *Col8a2*^Q455K/Q455K^ mice (**Figure 5d**). This was confirmed by Alizarin Red staining (**Figure 5e**), which demonstrated significantly higher corneal endothelium density in Ad-Cas9-Col8a2gRNA injected corneas than in uninjected FECD eyes (**Figure 5f**).

**Figure 5.**
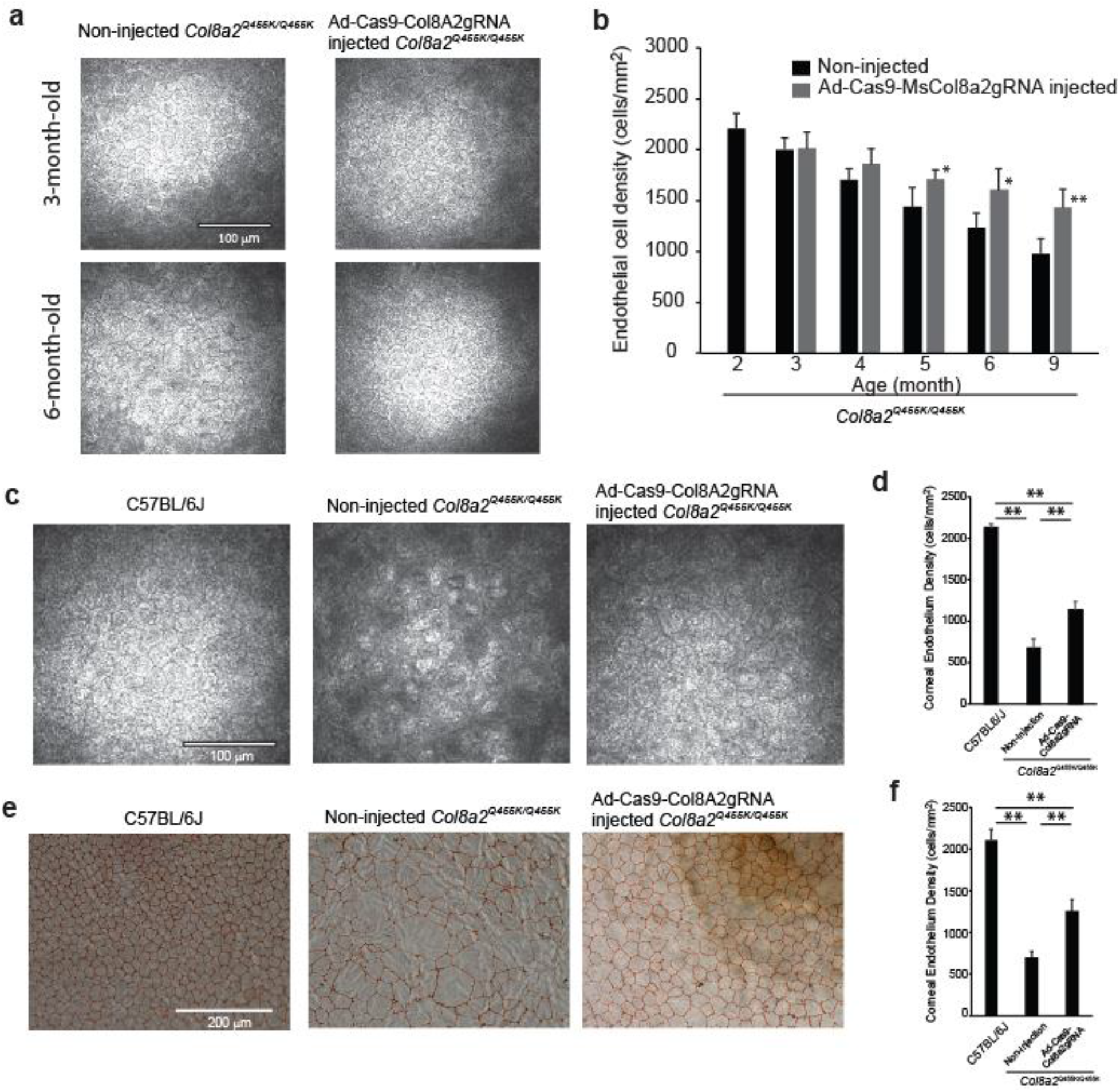
Ad-Cas9-Col8a2gRNA intracameral injection rescues corneal endothelium loss in the early-onset Fuchs’ dystrophy mice (*Col8a2*^Q455K/Q455K^) model. **a**, Representative *in vivo* corneal endothelium images using the Heidelberg Rostock microscope at 3- and 6-months post injection. Ad-Cas9-Col8a2gRNA was injected intracamerally into *Col8a2*^Q455K/Q455K^ mice at two months of age. Scale bar = 100μm. **b**, Time course change in corneal endothelial cell density of *Col8a2*^Q455K/Q455K^ mice, n=5. Ad-Cas9-Col8a2gRNA slows loss of corneal endothelial cells compared to no injection group. **c**, Representative *in vivo* corneal endothelium image at 12 months of age. Age-matched C57BL/6J and non-injected *Col8a2*^Q455K/Q455K^ mice were used for comparison. Ad-Cas9-Col8a2gRNA qualitatively improved endothelial cell density. Scale bar = 100μm. **d,** Average corneal endothelium densities: C57BL/6J: 2134±45 cells/mm^2^, non-injected *Col8a2*^Q455K/Q455K^: 677±110 cells/mm^2^ and Ad-Cas9-Col8a2gRNA injected *Col8a2*^Q455K/Q455K^: 1141±102 cells/mm^2^, n=4. Error bars show standard deviation. **e,** Representative corneal endothelium from each group stained with Alizarin Red. Scale bar = 200μm. **f,** Average corneal endothelium densities calculated from Alizarin Red stained corneas: C57BL/6J: 2108±134 cells/mm^2^, non-injected *Col8a2*^Q455K/Q455K^: 702±66 cells/mm^2^ and Ad-Cas9-Col8a2gRNA injected *Col8a2*^Q455K/Q455K^: 1256±135 cells/mm^2^, n=4. Error bars show standard deviation.

Further detailed analysis of corneal endothelium indicated changes in cell density and morphology (**Figure 6**). Analysis of paired corneas (injected and non-injected in the same mouse) showed significant improvement of corneal endothelial cell density by Ad-Cas9-Col8a2gRNA treatment in all four cases (**Figure 6a**). **Figure 6b** shows the distribution of corneal endothelial cell area. The morphology of the corneal endothelium, as monitored by hexagonality and coefficient of variation (COV) of its density were improved considerably (**Figure 6c-d**). *In vivo* corneal optical coherence tomography (OCT) demonstrated that Ad-Cas9-Col8a2gRNA decreased the formation of guttae-like structures compared to control (**Figure 7a-b**), which was confirmed by histology (**Figure 7c-d**). Thus, Ad-Cas9-Col8a2gRNA successfully ameliorated the loss of corneal endothelium and morphologic phenotype in the early onset FECD mouse model.

**Figure 6.**
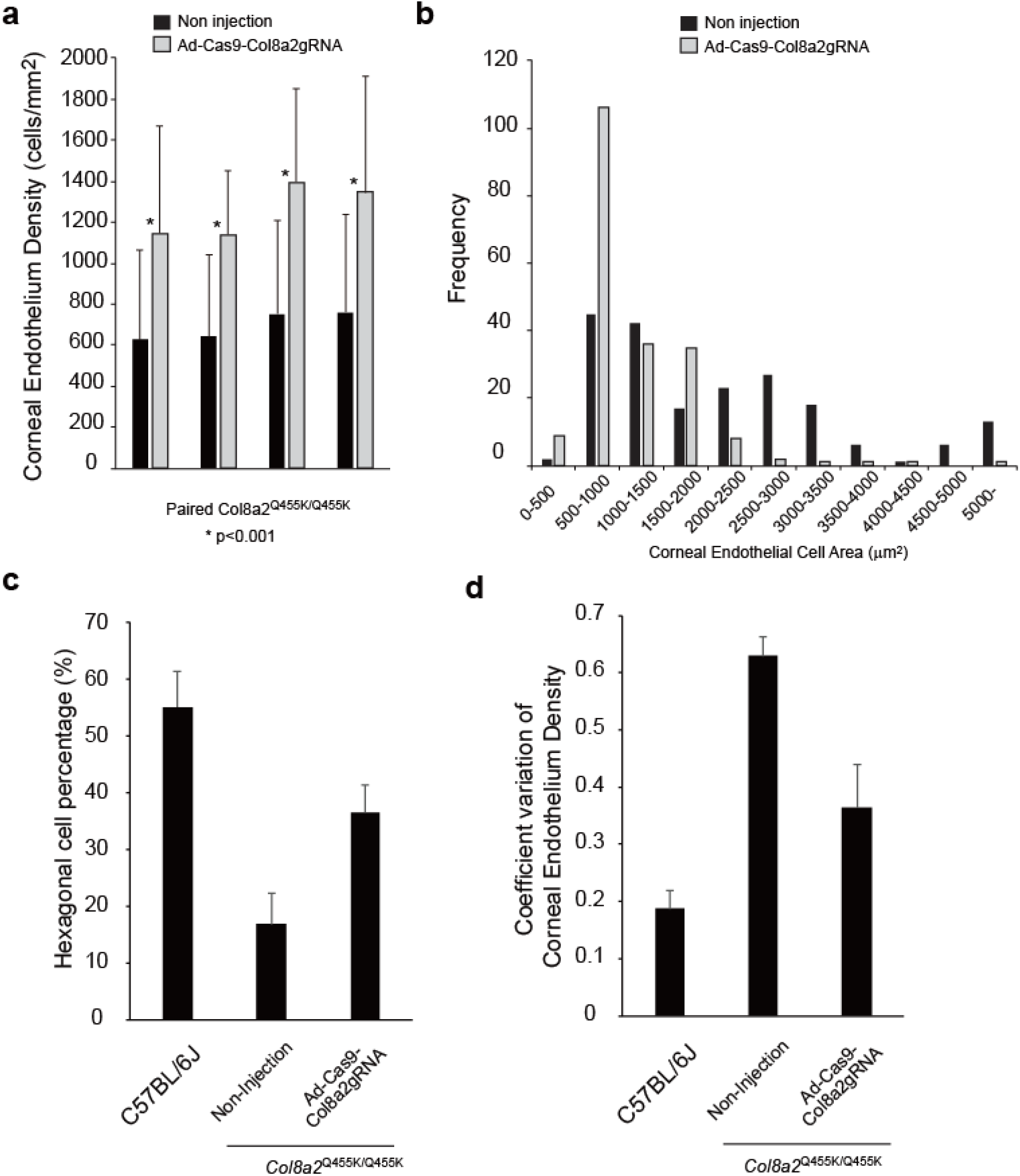
Ad-Cas9-Col8A2gRNA improves various characteristics of corneal endothelium in *Col8a2*^Q455K/Q455K^ mice. **a,** Corneal endothelium density in each cornea was calculated using Alizarin Red staining. A total of 50 different cell areas were measured in each cornea. Injected (Ad-Cas9-Col8a2gRNA) and uninjected corneas in the same mouse were compared by Student’s paired t-test. **b,** Histogram of corneal endothelial cell area in Ad-Cas9-Col8A2gRNA injected cornea and non-injected cornea quantitatively demonstrates left-shifting in cell size, i.e., enhanced density, in the former. N = 200 in each group from four different corneas. **c,** The hexagonality and **d,** coefficient of variation (COV) of corneal endothelium were significantly improved by Ad-Cas9-Col8A2gRNA intracameral injection in *Col8a2*^Q455K/Q455K^ mice. N =200 from four different corneas in each group.

**Figure 7.**
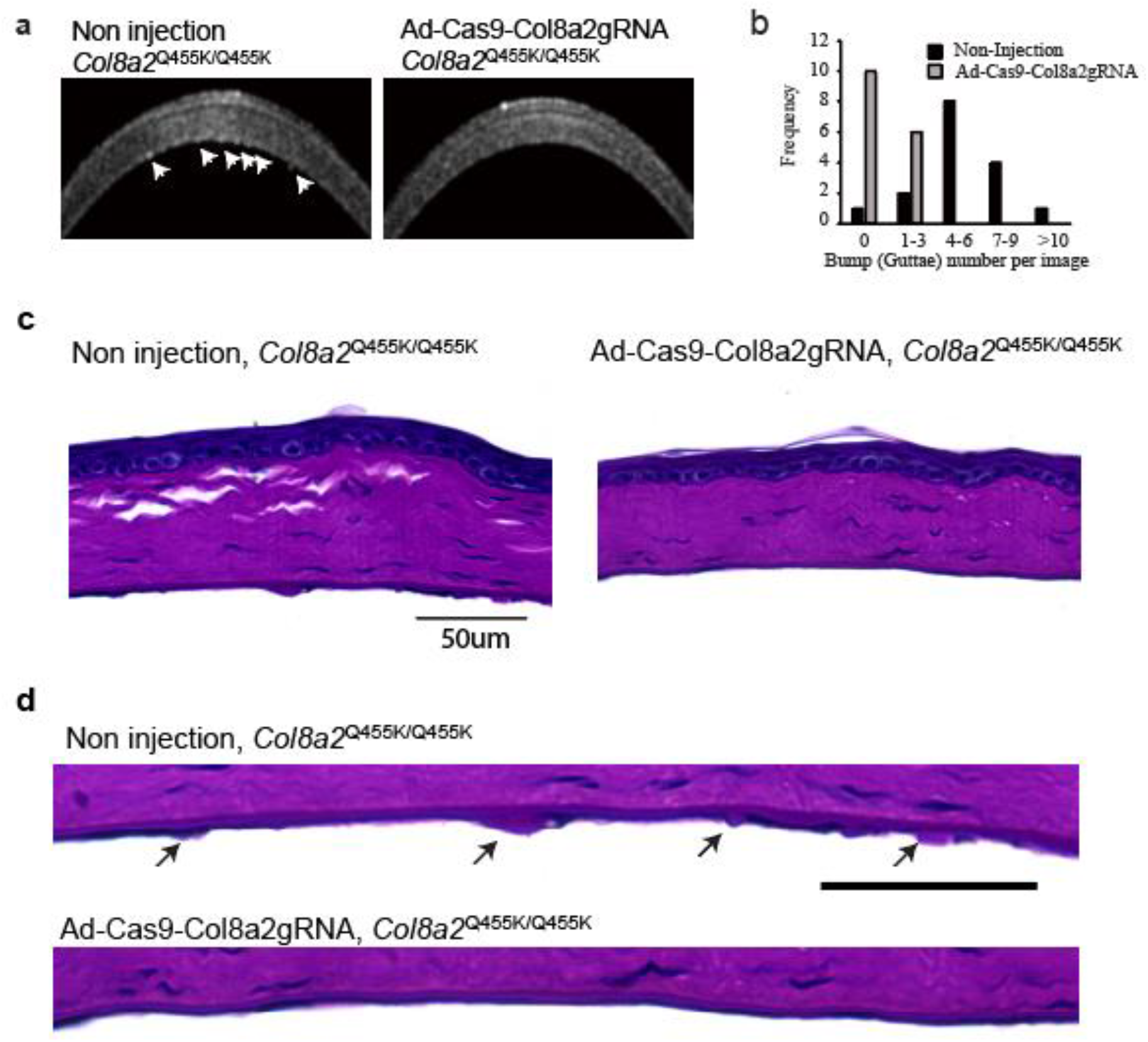
Ad-Cas9-Col8A2gRNA reduced guttae-like structures on the corneal endothelium in *Col8a2*^Q455K/Q455K^ mice. **a,** Corneal OCT revealed numerous guttae-like excrescences (arrows) in one year-old *Col8a2*^Q455K/Q455K^ mice, but far fewer in Ad-Cas9-Col8a2gRNA injected *Col8a2*^Q455K/Q455K^ mice. **b,** Histogram showing the number of guttae-like structures in each group. Non-injected *Col8a2*^Q455K/Q455K^: 5.2±3.4 excrescences/image and Ad-Cas9-Col8a2gRNA injected *Col8a2*^Q455K/Q455K^: 0.5±0.73 excrescences/image. n=16. P-value by Mann-Whitney U-test is <0.0001. **c-d,** PAS-stained corneas from uninjected and Ad-Cas9-Col8a2gRNA injected Col8A2^Q455K/Q455K^ mice. The arrows indicate guttae-like structures (excrescences).

### Ad-Cas9-Col8a2gRNA rescues corneal endothelium function in *Col8a2*^Q455K/Q455K^ FECD mice

Next, we examined whether Ad-Cas9-Col8a2gRNA could rescue corneal endothelial pump function of the *Col8a2*^Q455K/Q455K^ FECD mouse, which is essential for corneal clarity and optimal vision^23^. Surprisingly, *Col8a2*^Q455K/Q455K^ corneas do not develop edema or opacity even at one year of age despite reduced endothelial density (**Supplemental Figure 13**). We therefore developed a functional assay to deliberately induce corneal swelling and assess pump function by measuring the de-swelling rate. As direct application of a 0 mOsm/L solution was found to induce epithelial rather than stromal swelling (**Supplemental Figure 14**), we performed epithelial debridement to eliminate any confounding epithelial effects (**Figure 8a**). Application of an osmolar range of PBS solutions (**Figure 8b**) produced a range of swelling volumes, with 600-700mOsm/L solution producing the maximal effect, with quadrupling of the stromal thickness (**Figure 8b, c**). Thus, the epithelial layer functions as a barrier to maintain stromal thickness, whereas hypertonic solutions would seem to induce aqueous humor ingression into the cornea. Having optimized our model, we measured de-swelling rates following a 10-minute application of 650m Osm/L PBS. Successive corneal OCT images showed that the rate of de-swelling in non-injected *Col8a2*^Q455K/Q455K^ corneas was significantly delayed compared to C57BL/6J control corneas. In contrast, Ad-Cas9-Col8a2gRNA injected *Col8a2*^Q455K/Q455K^ corneas demonstrated de-swelling rates similar to C57BL/6J corneas (**Figure 8d, e**). Thus, Ad-Cas9-Col8a2gRNA rescued corneal endothelial function in FECD mice.

**Figure 8.**
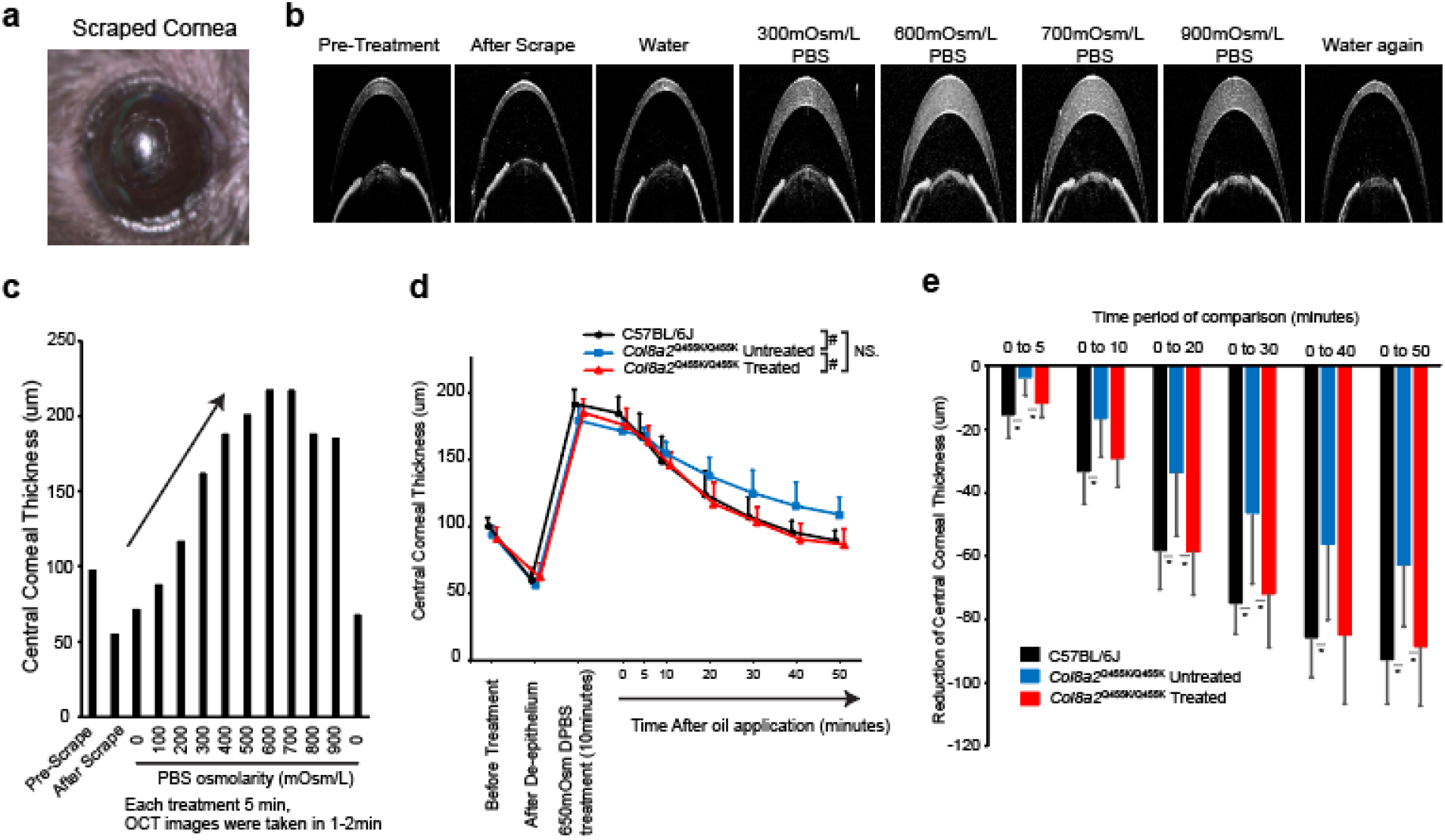
Ad-Cas9-Col8a2gRNA rescued corneal endothelium pumping function in *Col8a2*^Q455K/Q455K^ mouse. **a,** Stereomicroscopic images of scraped mouse cornea. **b,** Corneal OCT images of pre-treatment, after scrape, and after treatment with 0mOsm/L (water), 300, 600, 700, and 900mOsm/L DPBS application followed by water again. **c,** Changes in corneal thickness in response to variance inDPBS osmolality demonstrates that maximal swelling occurred at 600-700mOsm/L DPBS. **d,** Repeated measurements of central corneal thickness were taken using corneal OCT after application of 650mOsm/L PBS. To prevent evaporation, 4µL of silicone oil was applied at t = 0. (n=6). # indicated p<0.001 by regression analysis. NS: Not significant. **e,** Deswelling of central corneal thickness was measured from 0 min to 5, 10, 20, 30, 40, and 50 min. Non-injected *Col8a2*^Q455K/Q455K^ corneas showed significantly delayed deswelling compared to C57BL/6J corneas. In contrast, Ad-Cas9-Col8a2gRNA injection significantly improved corneal deswelling rate similar to that of C57BL/6J controls. (n=6). * indicated p<0.05 by Student’s t-test.

### Potential off-target of gRNA targeting the human *COL8A2* start codon

For potential therapeutic application of CRISPR/Cas9, we evaluated the off-target activity of humanized gRNA by a modified digenome analysis^24^. Briefly, digenome analysis consists of: 1) in vitro digestion of purified genomic DNA with SpCas9 and gRNA; 2) Deep sequencing of the digested genomic DNA; and 3) alignment of sequence reads at the digested sites. Consequently, digested sites other than the target site are considered potential off-target sites. In fact, we found the readings at the target site (human *COL8A2* start codon) were aligned but not without gRNA (**Figure 9a-b**). After careful observation, a gap was often found at the target site (**Figure 9c**). Since off-target analysis without considering such a gap would underestimate off-target events, we included a +/−1 gap in our modified digenome analysis. **Figure 9d** shows the digenome score alignments of control gDNA (no gRNA) and treated gDNA (HuCol8a2gRNA). From this, candidate sites were selected for which the score was >60. We identified 8 different sequences in 13 different locations that had homology to HuCol8a2gRNA and was associated with a PAM sequence (**Table 4**). The majority of these were non-coding sites, and the remaining sites (2 of which were anti-sense sites and 2 of which were intronic sites) (*SRGAP2-AS1, SV2c, KAT6B, LMO7-AS1, ACAN*) have no known corneal function. **Supplemental Table 1** shows 8 of 21 candidates which had neither homology to HuCol8a2gRNA nor PAM sequence. **Supplemental Table 2** shows 4 sequences in control gDNA.

**Figure 9.**
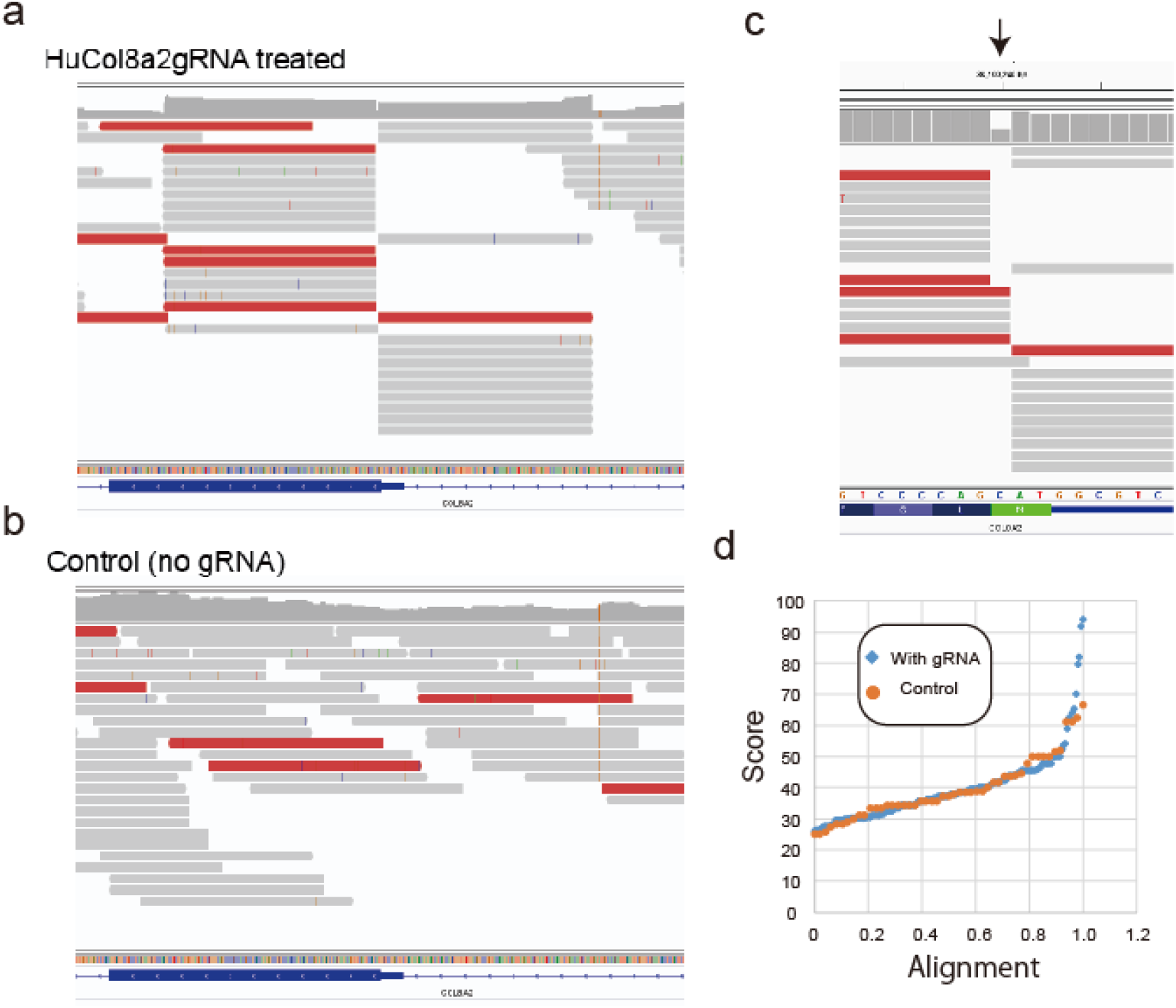
Modified digenome analysis for potential off-targets. **a, b,** Mapping of reads to human *COL8A2* target site from HuCol8a2gRNA treated gDNA and control gDNA. **c,** The gap was observed in *in vitro* digestion of genomic DNA. **d,** Modified digenome score alignment (0 to 1.0) of control gDNA (no gRNA) and HuGol8a2gRNA treated gDNA.

**Table 4.**
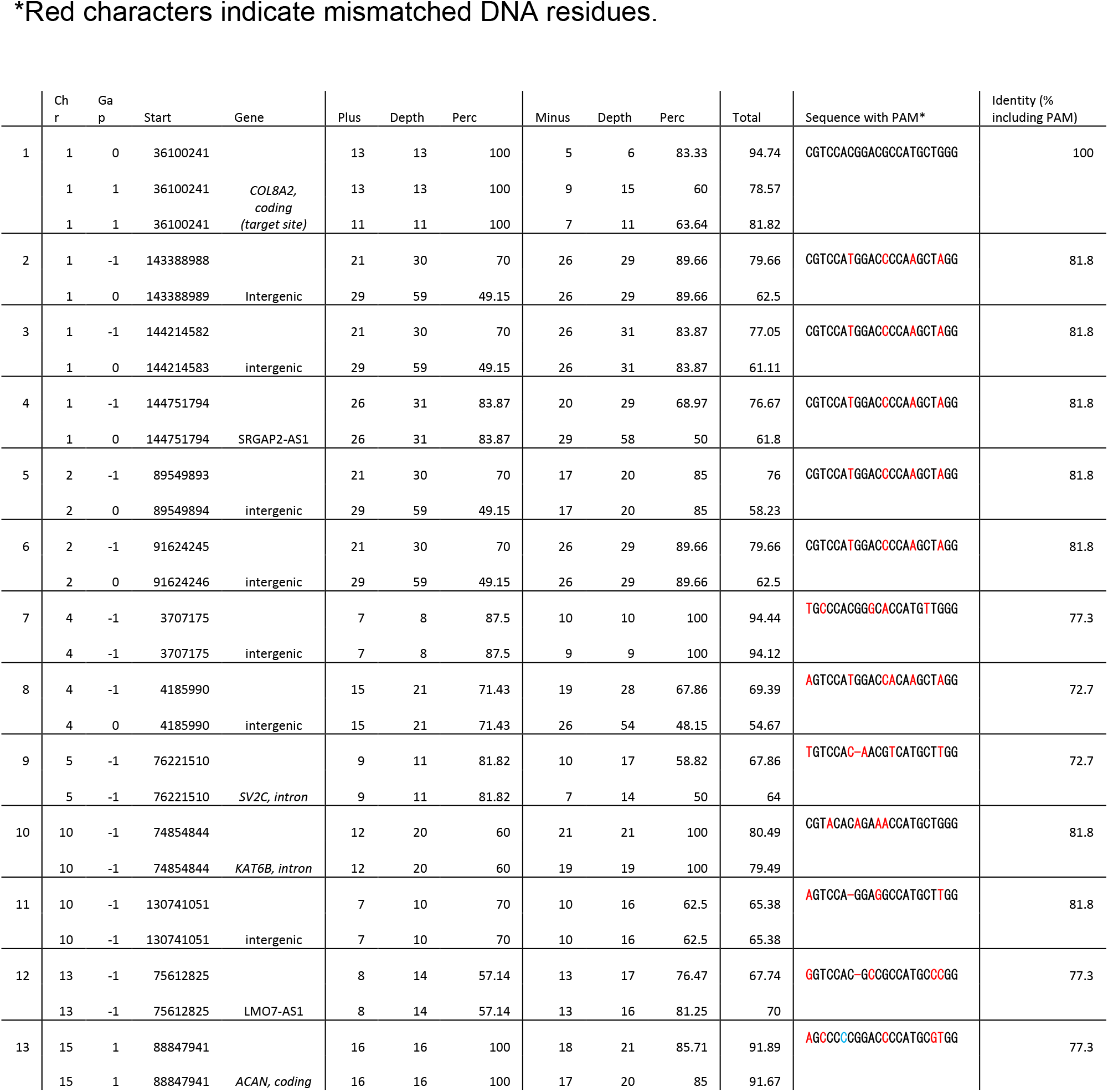
HuCol8a2gRNA off-target sites with homology.

## Discussion

In this study, we demonstrated that intraocular injection of a single adenoviral vector achieved efficient and restricted delivery of the CRISPR/Cas9 to adult post-mitotic corneal endothelium, leading to *in vivo* knock-down of mutant *Col8a2* with long-term preservation of corneal endothelial density, structure, and function in the early-onset Fuchs’ dystrophy mouse model.

We found that most of the insertions were single insertions of adenine, creating a cryptic start codon without frame shift (**Supplemental Figure 12**). As mentioned above, this would disrupt the kozak sequence. Taken together, our results indicate that disruption of the kozak sequence effectively reduces protein expression without complications such as non-functional or frame-shifted protein production. Hence, kozak sequence disruption by CRISPR/Cas9 targeting may provide a viable option for gene knock-down.

In this study, we performed the modified digenome method to determine potential off-target regions. Interestingly, we found a gap at the target site (**Figure 9**). This gap may have been generated during sample preparation, due to causes such as Covaris shearing, polishing of overhanging DNA and adenylation at 3’end for ligation or fluctuation of Cas9/gRNA recognition to genomic DNA. We identified 13 potential off-target sites with homology, the majority of which were in non-coding sequences and the other regions in genes of uncertain function. We found one potential coding exonic off-target sequence, in the *ACAN* gene. ACAN (also referred as aggrecan core protein) is a major component of extracellular matrix of cartilaginous tissues. Although several cartilage-bone related diseases are caused by mutations of *ACAN* coding region, its expression was not observed in previously published RNAseq data of human corneal endothelium^25,26^. Therefore, it is unlikely that this off-target indel affects corneal function. We found off-target sequences in an intron of two genes, *SV2C (Synaptic vesicle glycoprotein 2C)* and *KAT6B*. *SV2C* is involved in synaptic function throughout the brain^27^, but it it is rarely expressed in human corneal endothelium^25,26^. *KAT6B* is a histone acetyltransferase which may be involved in both positive and negative regulation of transcription. Several developmental disorders are caused by distinct mutations of *KAT6B*^28^, and acute myeloid leukemia may be caused by a chromosomal aberration involving *KAT6B* gene^29^. Therefore, KAT6B gene should considered a gene at risk with our Crispr/Cas9 treatment. In most cases, intronic mutations causing human diseases are located within 100bp from intron-exon boundary, as most diseases associated with intronic mutation create a pseudo-exon which disrupts splicing. The observed KAT6B off-target site is located over 11000bp from exon-intron boundary. Hence, the off-target mutation in *KAT6B* is unlikely to cause corneal dysfunction. Two additional off-target candidates were found in intron of non-coding RNAs, SRGAP2-AS1 and LMO7-AS1. Non-coding RNAs are sometimes known to have various functions in gene regulation, but the functions of SRGAP2-AS1 and LMO7-AS1 are unknown. All other off-target candidates are located in intergenic regions. Since some intergenic region contain gene enhancer element, mutations could theoretically contribute to disease risk^30^. Compared with exonic or intronic mutations, the risk of intergenic mutations inducing deleterious effects would be low. Thus, we identified off-target candidates of our Crispr/Cas9 treatment that would be expected to not cause corneal dysfunction. However, testing in large animals such as non-human primates should be performed prior to any clinical testing of in vivo crispr/cas9 treatment for humans.

Eight potential off-target sites without homology or PAM sequence were found, but we speculate these are likely random errors since the non-gRNA control also showed 4 potential off-target sites.

Previous papers have achieved *in vivo* editing in post-mitotic neurons using dual AAVs to co-infect cells with Cas9 machinery^31–33^. Although AAV has the advantages of low immunogenicity and toxicity, the low efficiency of homologous recombination by dual AAV delivery (10-12%)^31^ is unrealistic as a treatment approach, and the complexity of two vectors makes targeting efficacy assessment and clinical development challenging. Moreover, the long-term expression of AAV-based CRISPR/Cas9 may ultimately prove undesirable for a post-mitotic cell, since the potential for off-target gene editing will continue for the life of the AAV. In contrast, the high infectivity and short duration of adenoviral expression would enable structural and functional rescue by Ad-Cas9-Col8a2gRNA at a titer below adenoviral cytotoxicity, without risk of further (*mis*) editing events.

In conclusion, we succeeded in Col8a2 gene knock-down in corneal endothelium in vivo using an adenovirus mediated SpCas9 and gRNA delivery, resulting in a functionally relevant rescue of corneal endothelium in the early-onset FECD mouse model. Our strategy can be applicable to other genes and useful in experiments. However, prior to clinical development, gene therapy approaches will require optimization of gRNA and Cas9, understanding long-term effect, and refinement of the delivery strategy. Still, these results strongly suggest that our strategy can treat or at least prolong corneal endothelial life in early onset Fuchs’ dystrophy, potentially eliminating the need for transplantation.

## Materials and Methods

### Mice

C57BL/6J mice, 8-12 weeks old, were purchased from The Jackson Laboratory (Bar Harbor, ME) and used in this study. The *Col8a2*^Q455K/Q455K^ mouse has been previously described^17,18,34^. All animals were treated according to the ARVO Statement for the Use of Animals in Ophthalmic and Vision Research.

### Plasmid construction

px330 plasmid encoding humanized S. pyogenes Cas9 was obtained from Addgene (Cambridge, MA). The design of guide RNA (gRNA) and cloning were performed following published methods^20^. Three separate gRNAs were designed to target sequences containing a trinucleotide PAM sequence (in italics):

Col8A2-gRNA1: CCCATCCACAGACGCCATGC*AGG*;

Col8A2-gRNA2: GGGTGCAGCGGGCTATGCCC*CGG*;

Col8A2-gRNA3: CCGCCTTTCCGAGAGGGCAA*AGG*.

### Cell culture, plasmid transfection, and indel detection

Mouse NIH3T3 cells were obtained from ATCC (Manassas, VA) and maintained in 10% bovine calf serum/Dulbeccòs Modified Eaglès medium following manufacturer’s instructions. 2μg of plasmid was transfected by nucleofection (Lonza, Allendale, NJ). After two days, genomic DNA was purified using QIAamp DNA Mini Kit (Qiagen, Valencia, CA). 10ng of genomic DNA was PCR amplified with the following primer set; MsCol8a2_intron2F: cggtggtaggtggtaattgg and MsCol8a2_intron3R: tgtggtctggagtgtctgga. The PCR product (560bp) was purified with a Qiagen PCR purification kit and subsequently digested by CviAII restriction enzyme (NEB, Ipswich, MA) or Hin1II (Thermo Fisher Scientific, Waltham, MA) following the manufacturer’s protocols. We initially used CviAII before switching to Hin1ll due to low stability of CviAII (both enzymes cut CATG). Digested products were run on a 1% agarose electrophoresis gel. Uncut bands (~420bp) were purified and cloned with CloneJET PCR Cloning kit (Thermo Fisher Scientific). After transformation to DH5α (NEB), individual colonies were cultured in LB medium with ampicillin, purified via miniprep, and sent to the University of Utah DNA core facility for Sanger sequencing.

### Adenovirus production

Adenovirus production was carried out following previously published methods^35^. All restriction enzymes described here were purchased from NEB. Empty Shuttle vector (pShuttle, #16402) was obtained from Addgene. Col8a2-gRNA1 with U6 promoter and terminator was amplified from pCas9-Col8A2gRNA by PCR using the following primers; gRNAcloneF_EcoRV: TAGATATCgagggcctatttcccatgattc and gRNAcloneR_XbaI: TATCTAGAagccatttgtctgcagaattggc. PCR product was cloned into pShuttle using EcoRV/XbaI (pShuttle-Col8A2gRNA). Next, Cas9 DNA (including the promoter and polyadenylation signal) was excised from px330 with NotI/XbaI and cloned into pShuttle-Col8A2gRNA1 (pShuttle-Cas9-Col8A2gRNA1). After linearization with PmeI, pShuttle-Col8A2gRNA was electroporated into BJ5183-AD-1 cells (Agilent Technologies, Santa Clara, CA) and grown on kanamycin LB plates. Small colonies were individually picked and cultured in 5mL LB medium with kanamycin. After confirming size by digestion with PacI and other restriction enzymes, XL10-Gold Ultracompetent Cells (Agilent Technologies) were transformed with amplified plasmid of the correct size. The Maxiprep (Qiagen) purified plasmids were linearized by PacI digestion, and transfected to AD-293 cells (Agilent Technologies) using Lipofectamine® 2000 (Thermo Fisher Scientific). After 14-20 days culture, adenovirus generating AD293 cells were harvested. HeLa cells were used to confirm the replication deficiency. The titer of recombinant adenovirus was determined by Adenovirus Functional Titer Immunoassay Kit (Cell Biolabs, Inc., San Diego, CA). The function of Ad-Cas9-Col8a2gRNA was examined using NIH3T3 as described above. For *in vivo* experiments, further production and purification were performed in a viral core facility at the University of Massachusetts.

### Anterior chamber injection

Eight-week-old male C57BL/6J mice received a single unilateral injection of Ad-Cas9-Col8a2gRNA into the anterior chamber, while the contralateral eye served as an uninjected control. All injections were performed in an Animal Biosafety Level 2 Comparative Medicine Core Facility at the University of Utah. Mice were first anesthetized with ketamine (90mg/kg) and xylazine (10mg/kg) before topical application of tropicamide and proparacaine. Corneas were punctured 1.5mm above the limbus with a 31G needle and the needle gently withdrawn. Using a blunt 33G Hamilton syringe, Ad-Cas9-Col8a2gRNA (4μL) was injected through the puncture. To ensure injection delivery, the cannula remained in the anterior chamber for ~5 seconds after injection before applying erythromycin ophthalmic ointment to the cornea.

### Measurement of indel rate by deep sequencing

One-month post Ad-Cas9-MsCol8a2gRNA to C57BL/6J mice, the corneal endothelium was separated mechanically (**Supplemental Figure 2**). Genomic DNA from the corneal endothelium/stroma was purified by Quick-DNA Microprep Plus Kit (Zymo research). PCRs were performed on the locus using TTCTTCTTCTCCCTGCAGCC and GCACATACTTTACCGGGGCA (30 cycles, the product size: 155bp). The deep sequencing was performed by the HSC core at University of Utah. The library was prepared using the Swift Biosciences Accel-NGS 1S Plus DNA Library Kit. The sequence protocol used MiSeq Nano 150 Cycle Paired End Sequencing v2. The total number of reads per file was counted. The reads with median quality score ≤5 were removed from the data set. The reads were aligned to the expected genomic sequence: gi|372099106|ref|NC_000070.6|:126309560-126309770 Mus musculus strain C57BL/6J chromosome 4, GRCm38.p4 C57BL/6J.

### Digenome sequencing

a. **Human *COL8a2* gRNA design** We designed two different human col8a2 gRNAs at the start codon of human *COL8A2* similar to mouse *Col8a2* gRNA. HuCol8a2gRNA1 ACGTCCACGGACGCC<u>ATG</u>C HuCol8a2gRNA2 CGTCCACGGACGCC<u>ATG</u>CT Underlines indicate the start codon of human *COL8A2*. As explained in the main text, these sequences were cloned into px330 plasmid.
b. **AD-293 cell culture and plasmid transfection** To confirm the activity of human gRNAs, we used human AD-293 cells (Stratagene), which were maintained following the manufacturès instructions. Ca-phosphate method was used for plasmid transfection. Briefly, 0.25 × 10^6^ cells were plated in 6-well plate with 2mL of 10%FBS/DMEM. Next day, 6µg plasmid were transfected. Two days post transfection, genomic DNAs were purified with Quick-DNA Plus Kit (ZymoResearch).
c. **PCR and restriction enzyme digestion for indel examination** To examine the indel at the target site, we used PCR and restriction enzyme digestion. PCR primers are; HuCol8a2_F: tgatcttttggtgaccccgg, HuCol8a2_R: GGATGTACTTCACTGGGGCA. The PCR product (226bp) was digested with Hin1II which recognizes CATG. Without indels, the *COL8A2* PCR products were digested to 94bp and 132bp. As shown in **Supplemental Figure 15a**, both px330 plasmids transfection showed the indel. Since we found HuCol8a2gRNA2 showed slightly higher activity, we proceeded with HuCol8a2gRNA2 for further experiments (Mentioned as HuCol8a2gRNA hereon).
d. **gRNA production by in vitro transcription** To produce gRNA, in vitro transcription was performed with MEGAshortscript™ T7 Transcription Kit (Thermofisher). The template DNA was obtained by PCR (Phusion® High-Fidelity DNA Polymerase, NEB) with primers (Forward: <u>TAATACGACTCACTATAG</u>CGTCCACGGACGCCATG, Reverse: AAAAGCACCGACTCGGTGCCA. The underline indicates T7 promoter.) using px330-huCol8a2gRNA plasmid as a template. The integrity of guide RNA was confirmed by 2% agarose DNA electrophoresis (**Supplemental Figure 15b**).
e. **In vitro genome digestion with Cas9** SpCas9 protein was obtained from NEB (M0386M). The reaction was performed in 8µg genomic DNA (AD-293), 120pmol (300nM) SpCas9, 120pmol (300nM) or 360pmol (900nM) gRNA with 1X NEBuffer™ 3.1 (total volume is 400µL) at 37℃ for 8 hours. After genomic DNA purification, the digestion at the target site was examined by PCR with HuCol8a2_F and HuCol8a2_R primers (**Supplemental Figure 15c**). We found that 360pmol gRNA (Cas9 : gRNA = 1 : 3) showed efficient digestion. Therefore, we proceeded with 360pmol gRNA treated gDNA for deep sequencing.
f. **Deep sequencing** Deep sequencing was performed at the HSC core at University of Utah. The library was prepared with Illumina TruSeq Nano DNA Sample Prep kit. The sequence protocol is NovaSeq 2 × 150 bp Sequencing 30X Human Whole Genome.
g. **Data analysis** The sequencing data was analyzed at the Bioinformatics core of the University of Utah. As shown in **Figure 9a-b**, Cas9-digested gDNA with HuCol8a2 gRNA showed aligned sequencing at the Col8a2 gene target site. On the other hand, control gDNA (Cas9-digested without gRNA) showed random sequencing. Since we found a gap at the target site (**Figure 9c**), our analysis accepts the gap which is explained below. The human GRCh38 FASTA file was downloaded from Ensembl and a reference database was created using bowtie2 version 2.3.4. Adapters were trimmed out of reads using Cutadapt 1.16 and then aligned using Bowtie 2 in end-to-end mode (full options -- end-to-end --sensitive --no-unal -k 20). The aligned reads were loaded into R using the GenomicAlignments package and total coverage and read start coverage were calculated for the plus and minus strands. Positions with 5 or more read starts were compared to the total coverage and read starts with less than 25% of total coverage were removed. The filtered read starts on the positive and negative strands were joined to find predicted cut sites with either no overlap (blunt end), 1 base pair gap or 1 base pair overhang.

### *In vivo* optical coherence tomography and corneal confocal microscopy

Two months after anterior chamber injection, corneal thickness was quantified by Spectralis OCT with the anterior-segment OCT module (Heidelberg Engineering, Franklin, MA). A HRT3 Rostock microscope (Heidelberg Engineering) was used to produce serial images of central corneal endothelial density, and endothelial cell counts were performed using ImageJ.

### Immunohistochemistry and histology

Immediately following mouse euthanasia, eyes were enucleated and the sclera/retina was punctured to facilitate fixation by immersion in 4% paraformaldehyde/PBS at 4℃. After 2 hours of fixation, the cornea was excised at the limbal boundary, paraffin embedded using standard protocols, and sectioned at 10μm. For COL8A2 immunostaining, we used avidin-biotin based detection (Vector Lab *Elite* ABC kit, Burlingame, CA) with 5µg/ml rabbit anti-COL8A2 polyclonal antibody (PA5-35077, Thermo Fisher Scientific). 5µg/ml rabbit IgG, was used as an isotype control (02-6102, Thermo Fisher Scientific). After developing with DAB (Vector Lab), and counter-staining with Nuclear Fast Red (Vector Lab), 20x magnified images were obtained with a light microscope (EVOS FL Auto Cell Imaging System, Thermo Fisher Scientific). Masson’s trichrome and Periodic Acid-Schiff (PAS) staining were performed using Trichrome Stain Kit (Masson, HT15, Sigma-Aldrich, St. Louis, MO) and PAS Kit (395B, Sigma-Aldrich) respectively. For corneal endothelial cell density, the whole cornea was fixed with acetone for 1 hour. This and all subsequent washes and incubations were performed at room temperature. After 4 washes with PBS, the cornea was blocked for 1 hour (3% BSA/PBS) and incubated for a further hour with 2.5µg/ml Alexa Fluor® 488 conjugated to anti-ZO1 antibody (339188, Thermo Fisher Scientific). After four final PBS washes, corneas were mounted on glass slides, endothelial side up, and imaged by confocal microscopy (Olympus FluoView FV1000). Corneal endothelial density was calculated manually by counting the number of corneal endothelial cells in three different areas of each cornea.

For immunostaining on corneal cryosections, we used rat anti-TNFα antibody (clone MP6-XT22, BioLegend, San Diego, CA) and rat anti-IFNγ (clone XMG1.2, BioLegend). As a control, we used isotype antibody (RTK2071, BioLegend). Briefly, the sections were blocked with 5% goat serum, 0.02% triton X-100/PBS for 30 minutes at room temperature. Then, the sections were stained with each antibodies at 5mg/mL for 1 hour at room temperature. After washing with PBS, the sections were stained with Alexa Fluor 647 conjugated goat anti-rat IgG (H+L) antibody (A-21247, Thermo Fisher Scientific). After DAPI staining, the fluorescence was observed with EVOS microscope.

### Electroretinography

C57BL6J mice were injected with Ad-GFP (anterior chamber injection), Ad-Cas9-Col8a2gRNA (anterior chamber injection) or 1 μg concanavalin A (intravitreal injection) (sigma). The mice were examined with ERG for retinal function safety 0 (prior to injection), 2 and 4 Mice were dark-adapted overnight before the experiments and anesthetized with intraperitoneal injection of Tribromoethanol and 2-methyl-2-butanol diluted in physiological saline at 14,5mL/kg dose. The pupils were dilated with tropicamide (0.5%) and phenylephrine (2,5%) eye drops. ERG experiments were performed with a Ganzfeld ERG (Phoenix laboratories). Scotopic combined response was obtained under dark-adapted conditions (no background illumination, 0 cd/m^2^) using white-flash stimuli ranged from -1.7 to 1.0 log cd s/m^2^ with twenty responses averaged for each stimulus.

### Alizarin Red staining

Alizarin Red staining for corneal endothelium was performed according to previously published methods^36^. After euthanizing mice, corneas were harvested and washed twice with saline (0.9% NaCl) prior to a 2-minute immersion in 0.2% Alizarin Red solution (pH 4.2 adjusted by 0.1% NH_4_OH, in saline). After washing twice again with saline, corneas were fixed with acetone for 10 minutes and again washed in saline three times (10 minutes each). Corneas were mounted on glass slides and imaged with a bright field microscope.

### Corneal Swelling/De-swelling experiment

Mice were anesthetized with ketamine/xylazine. Imaged corneas were kept moist with DPBS, excess DPBS was removed with absorbent tissue, while the contralateral eye was covered with ointment to prevent dehydration. Corneal OCT images were taken before scraping and before treatment. The corneal epithelium was removed mechanically using a Tooke corneal knife (Novo Surgical Inc., Oak Brook, IL) and jeweler’s forceps (**Figure 8a**). This process takes about 5 minutes. For testing the corneal swelling response to different osmolalities of DPBS solution, we sequentially applied solutions at 5-minute intervals, beginning with 0mOsm/L (deionized water) to 900mOsm/L DPBS, completely covering the eye throughout the course of each application. Each application required 1-2 minutes for image acquisition with OCT, which was performed immediately after removing residual solution with clean absorbent paper. To analyze corneal de-swelling, the cornea was fully covered with 650mOsm/L DPBS for 10 minutes. After removing excess solution with clean filter paper, 4µL of silicone oil was applied to avoid evaporation from the corneal surface. Corneal and OCT images commenced at 5, 10, 20, 30, 40 and 50 minutes.

### Statistical Analysis

Student’s t-test was used for comparison of average accompanied with ANOVA for multiple group comparison. To compare the slopes of central corneal thickness trajectory, we employed linear mixed-effects regression approach among groups of C57BL/6J, Non-injected Col8a2^Q455K/Q455K^ and Ad-Cas9-Col8a2gRNA injected Col8a2^Q455K/Q455K^ mice. Random-effect component in the regression approach was used to account for the correlation among repeated measurements within each mouse. The regression analyses were performed using statistical software R at a significance level of 0.05.

## Acknowledgement

This work was supported by the National Institutes of Health / National Eye Institute (R01EY017950), a NIH/NEI core grant and an unrestricted grant from Research to Prevent Blindness, Inc. New York, NY. to the Department of Ophthalmology & Visual Sciences, University of Utah.

---

**Supplemental Figure 1.**
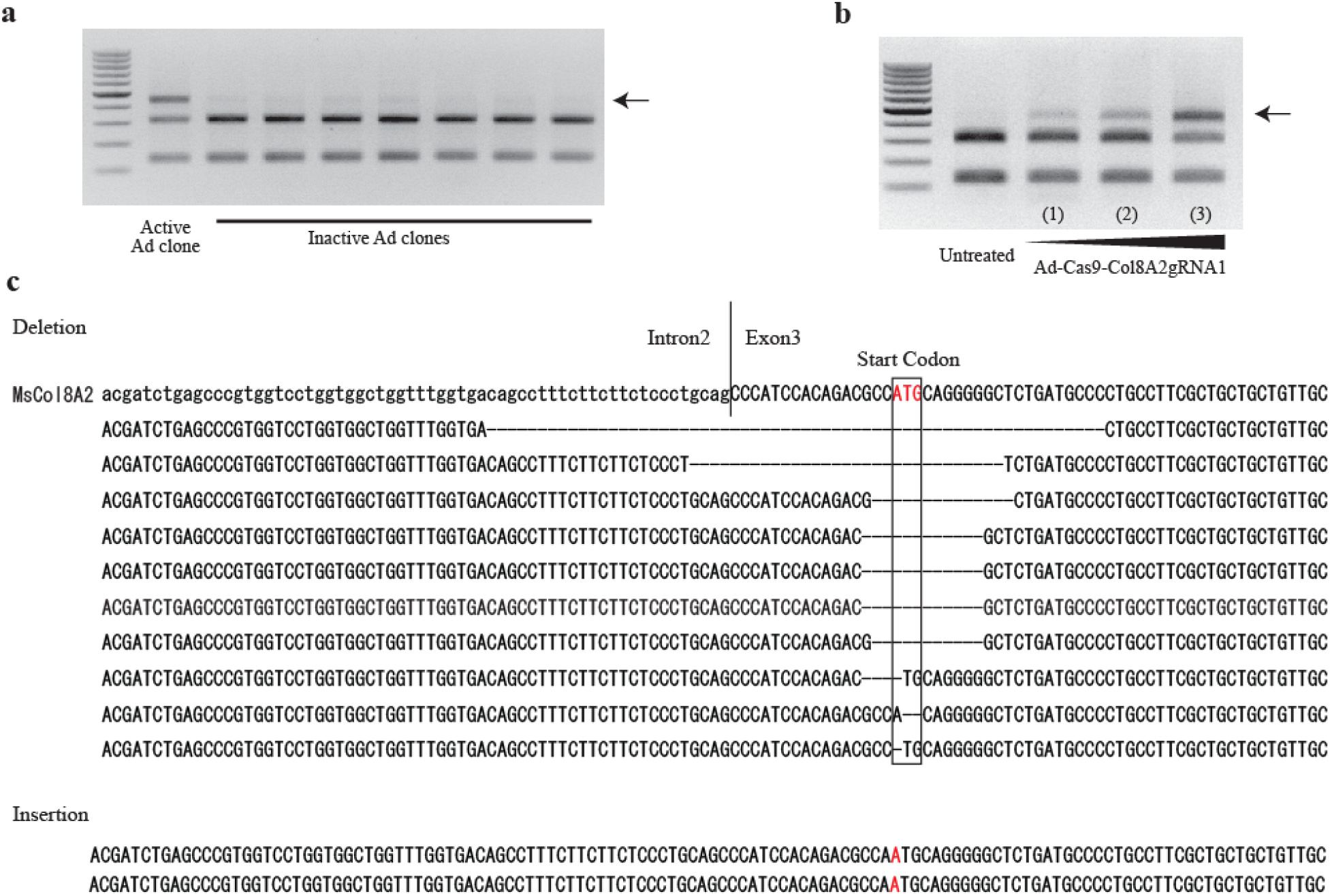
Ad-Cas9-Col8a2gRNA cloning and its indel activity *in vitro*. **a,** Cloning of Ad-Cas9-Col8a2gRNA by its indel activity in HEK293 cells. The method was the same as described for Figure 1 and supplemental Figure 1. The extra band (arrow) demonstrates the indel at the start codon. We examined 30 different clones and found only one clone with indel activity. **b,** As Ad-Cas9-Col8a2gRNA titer increased, indel activity also increased. **c,** Results from Sanger sequencing of cloned PCR products.

**Supplemental Figure 2.**
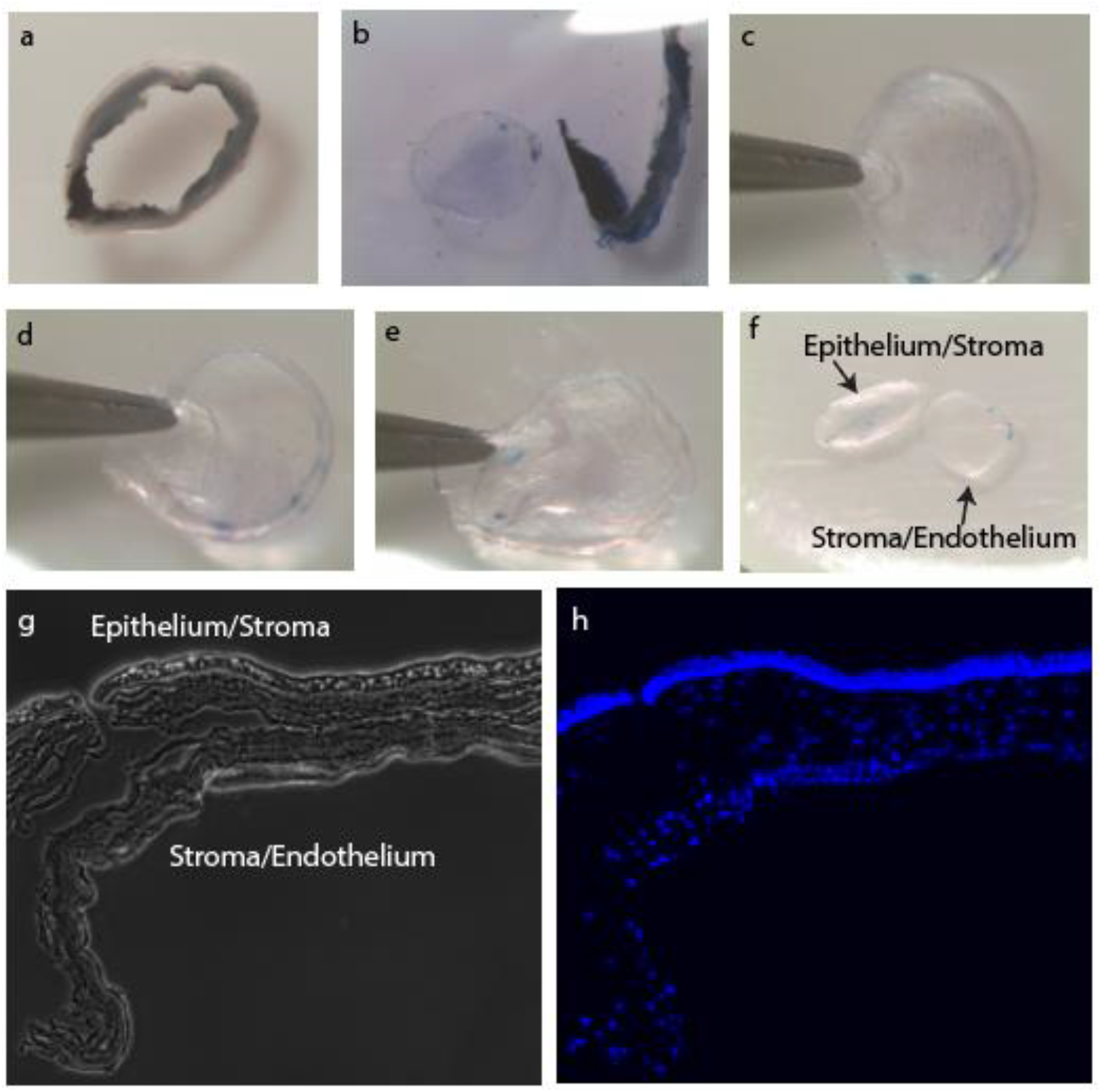
Procedure of peeling-off mouse corneal endothelium. Mouse corneal endothelium was peeled off mechanically. **a.** Mouse cornea after excision from of the rest of the eye. **b.** Mouse cornea was stained with 0.4% trypan blue for visualization, and the limbus/sclera was removed. **c-e.** Mechanical peeling of corneal endothelium. **f**, epithelium/stroma and stroma/endothelium after complete separation. **g-h,** Cryosection image of the cornea with endothelium peeled. DAPI staining showed incomplete separation of corneal endothelium and stroma.

**Supplemental Figure 3.**
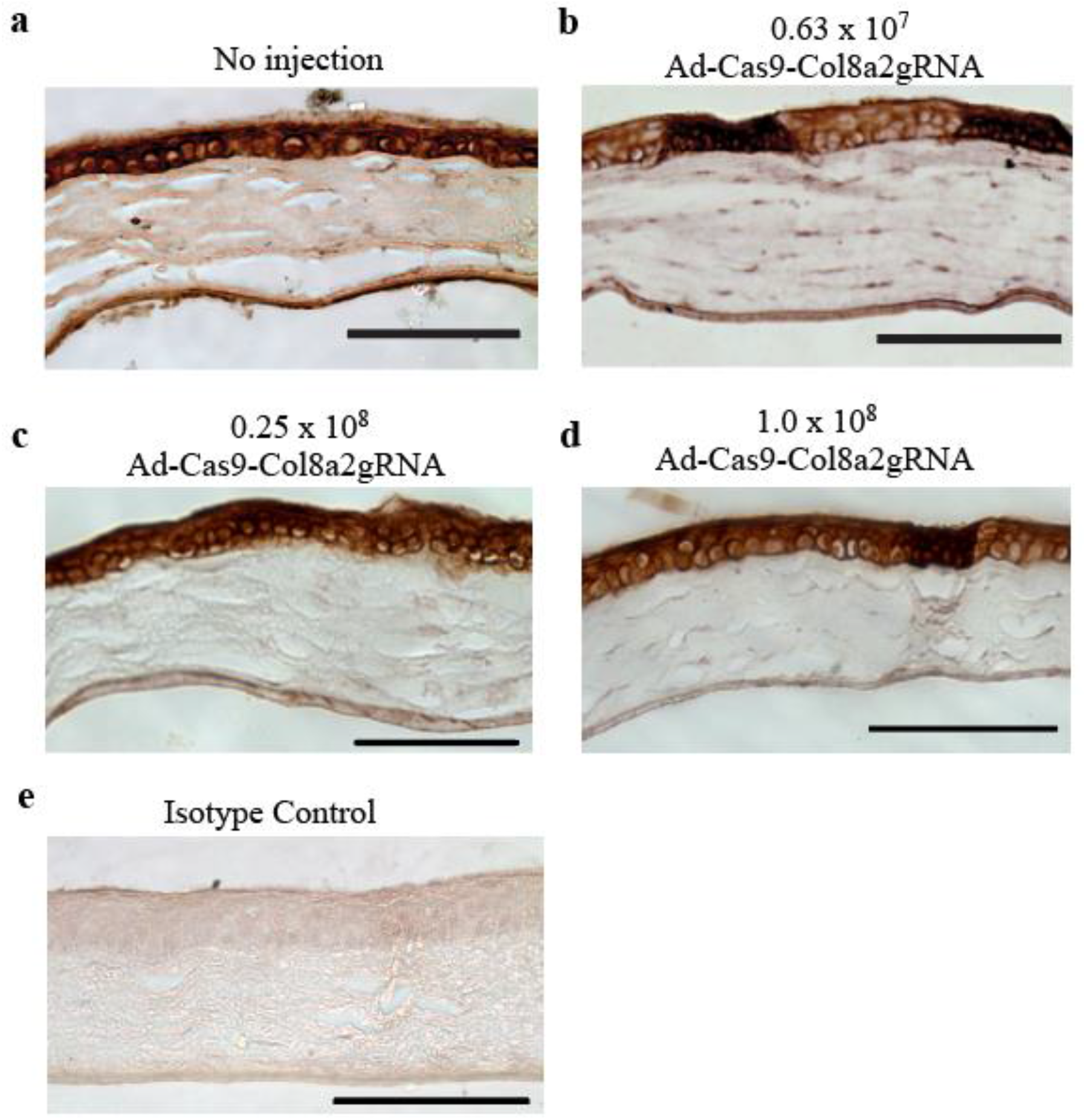
Various doses of Ad-Cas9-Col8a2gRNA reduce COL8A2 expression in C57BL/6J cornea. **a,** No injection. **b-d,** 0.63 × 10^7^, 0.25 × 10^8^ and 1.0 × 10^8^vg of Ad-Cas9-Col8a2gRNA in 4µL were injected intracamerally. Corneas were harvested two months post-injection. **e,** Isotype control. Scale bar is 100µm.

**Supplemental Figure 4.**
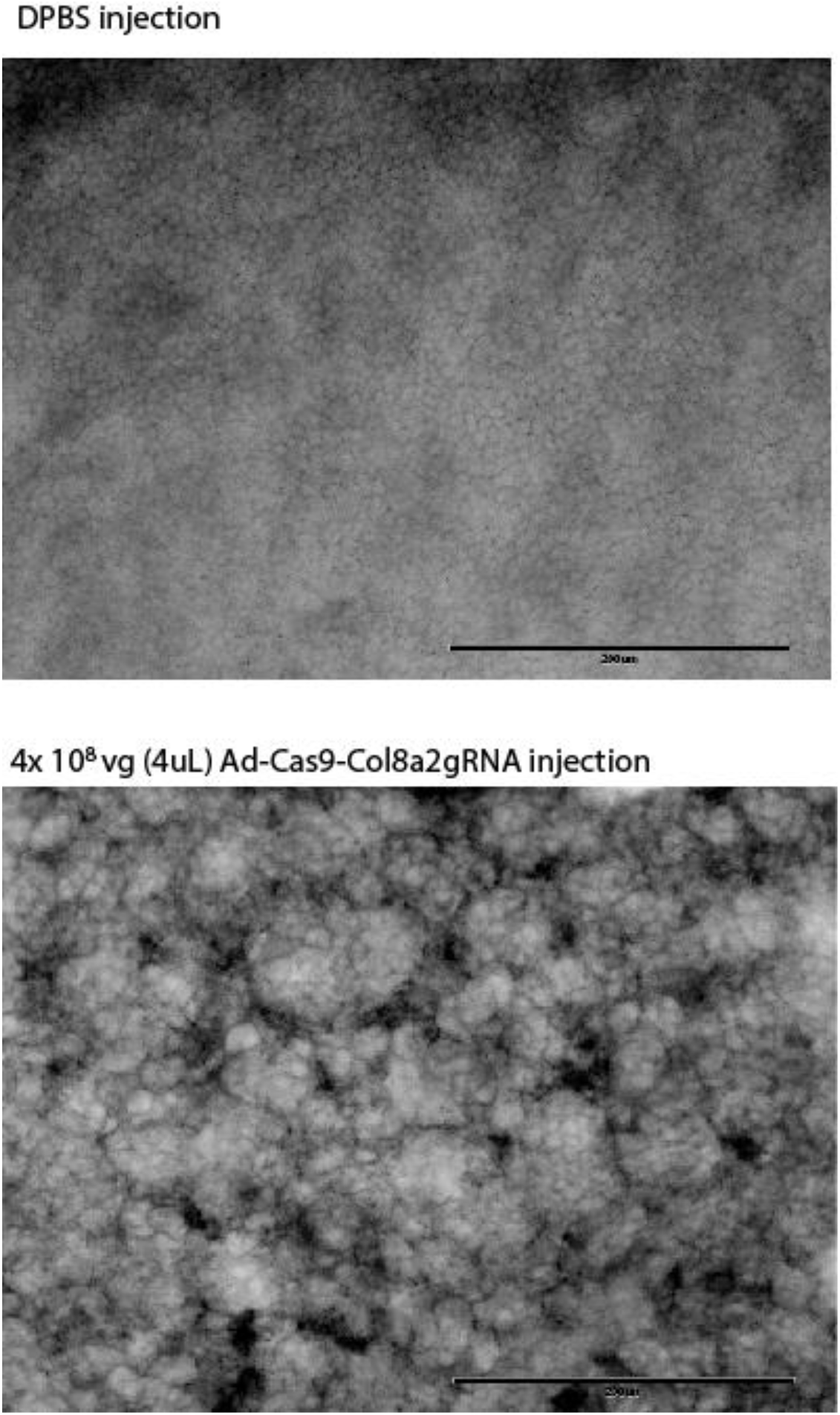
High level of Ad-Cas9-Col8a2gRNA (4 × 10^8^) is toxic to corneal endothelium in C57BL/6J mice. Two weeks following intracameral injection of DPBS or Ad-Cas9-Col8a2gRNA (4 × 10^8^vg), corneas were harvested to examine endothelial integrity with anti-ZO-1 antibody. This high titer of Ad-Cas9-Col8a2gRNA led to widespread devastation of the corneal endothelium. Scale bar = 200µm.

**Supplemental Figure 5.**
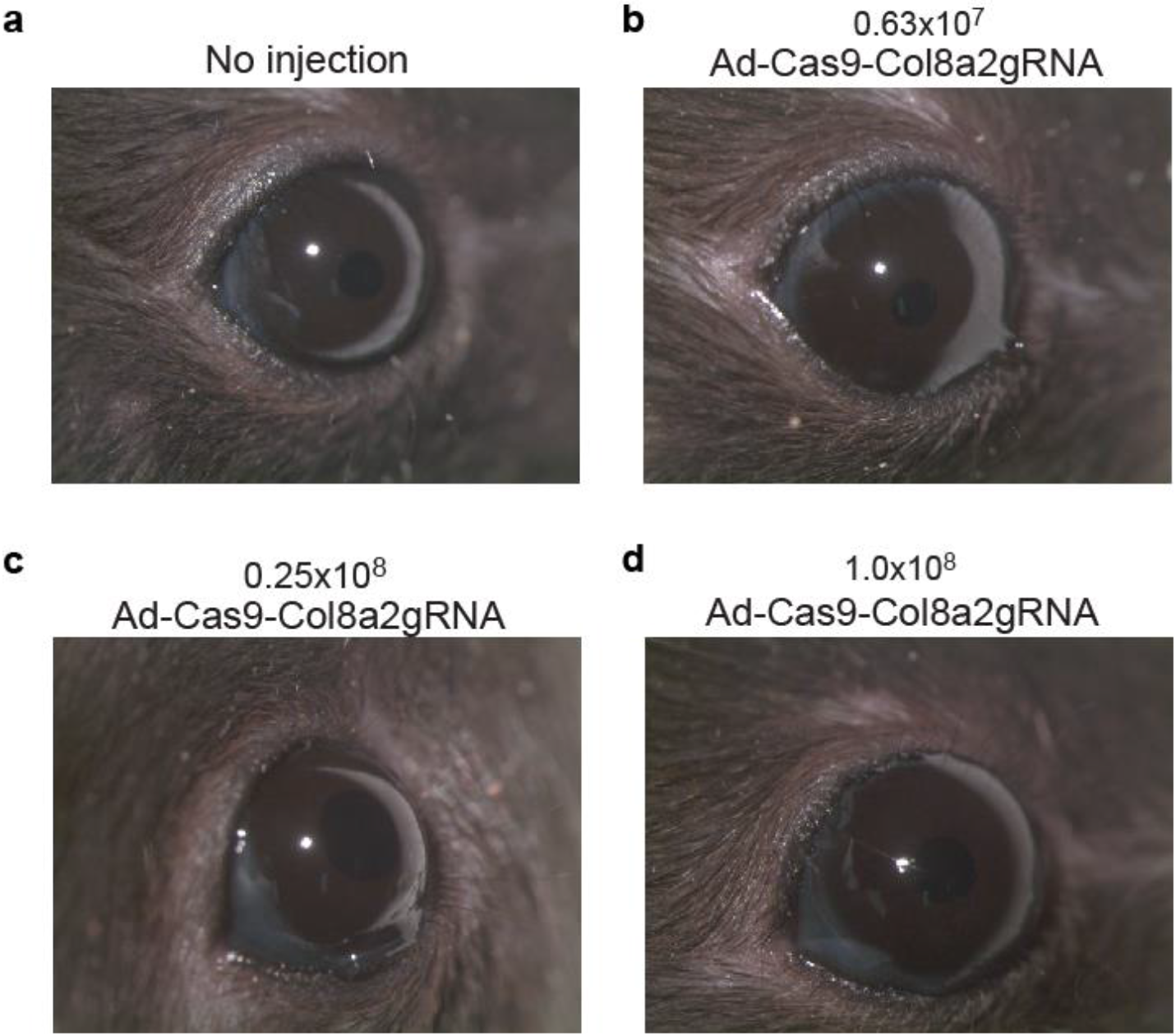
Low doses of Ad-Cas9-Col8a2gRNA intracameral injection did not induce corneal edema or clouding. **a-d,** Injection of Ad-Cas9-Col8a2gRNA at 0.63 × 10^7^, 0.25 × 10^8^ and 1.0 × 10^8^vg), did not result in corneal edema or opacity.

**Supplemental Figure 6.**
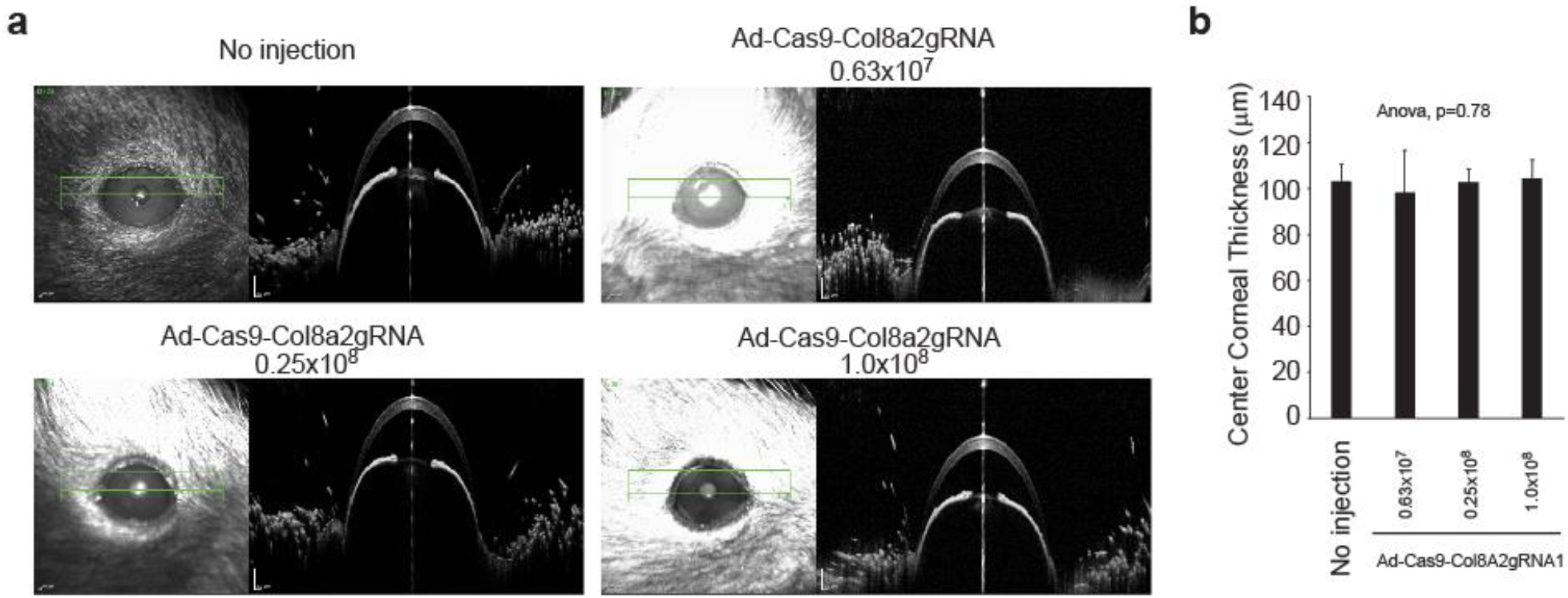
Baseline central corneal thickness is not different after injection of different titers of Ad-Cas9-Col8a2gRNA in C57BL/6J. **a,** Representative corneal OCT images captured by Heidelberg Spectralis microscope for each condition. **b,** The average of central corneal thickness in each condition. Significant differences among groups were not observed (ANOVA, p=0.78). n=6-8. Error bars show standard deviation.

**Supplemental Figure 7.**
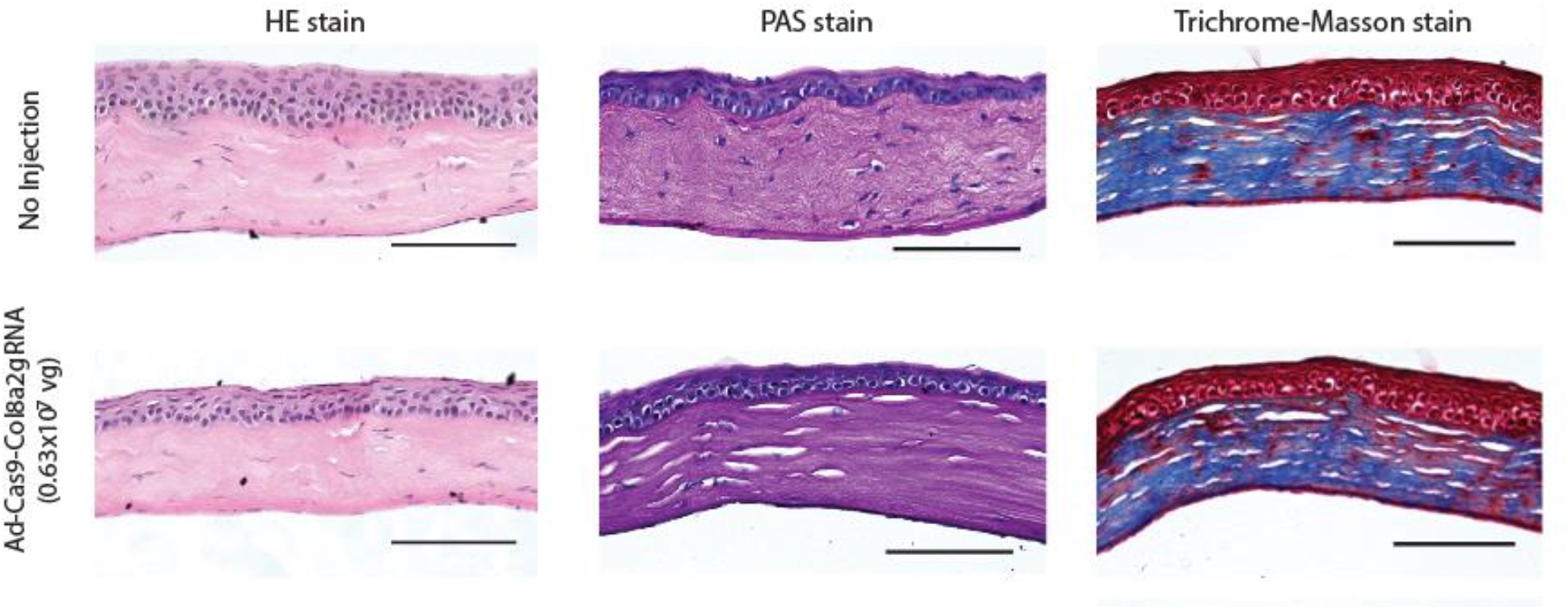
H&E, PAS, and Trichrome Masson staining showed no apparent phenotypes in Ad-Cas9-Col8A2gRNA injected corneas compared to uninjected corneas. There was no evidence of necrosis, inflammation, fibrosis, or other histologic changes to corneal architecture. Scale bar = 50µm.

**Supplemental Figure 8.**
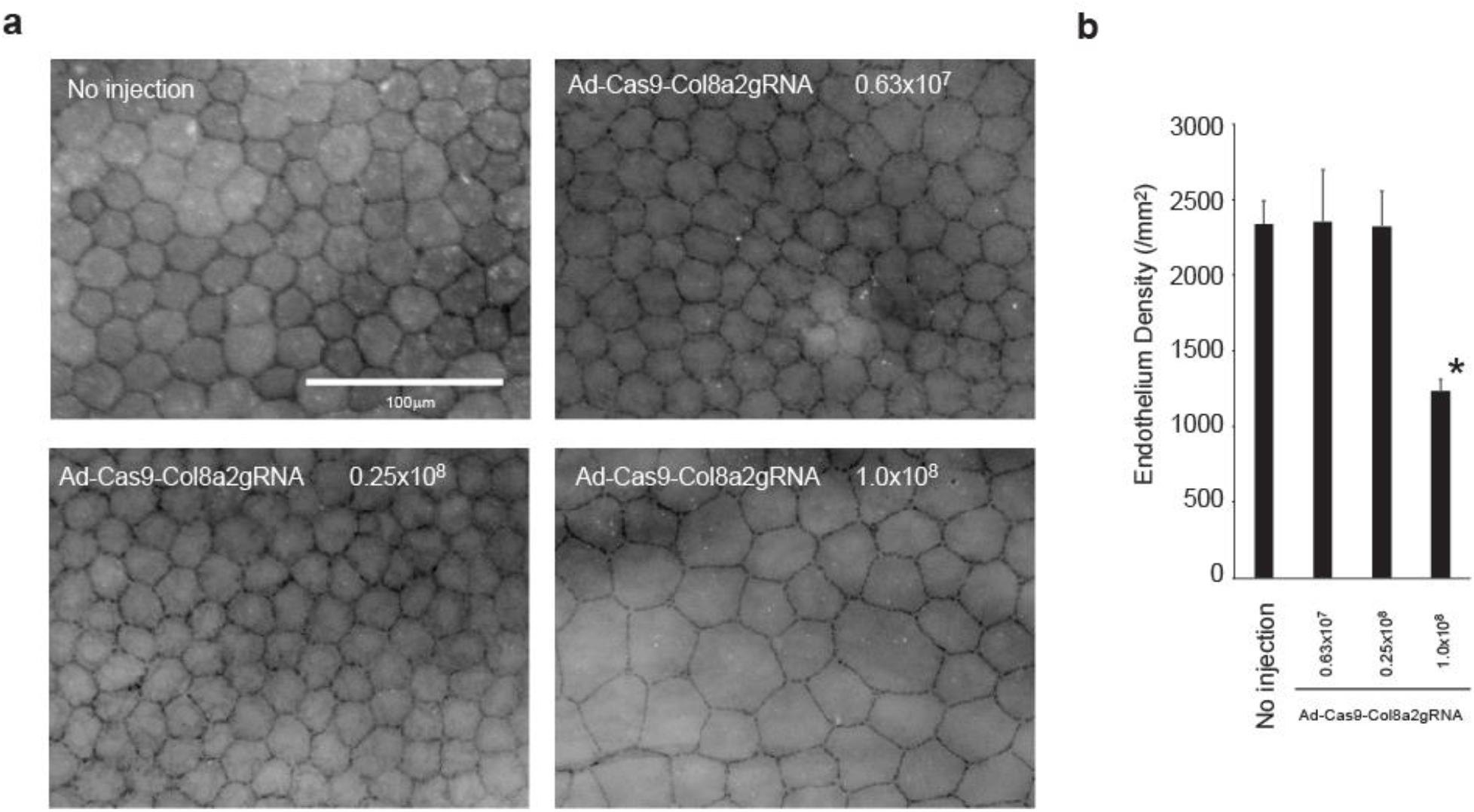
Intracameral injection of 1.0 × 10^8^ Ad-Cas9-Col8A2gRNA reduced corneal endothelium density in C57BL/6J mice. **a,** Representative images of corneal flat mounts immunolabeled with ZO-1 for each condition. Scale bar = 100µm. **b,** Average corneal endothelium densities. 1.0 × 10^8^vg Ad-Cas9-Col8A2gRNA reduced corneal endothelium density significantly. n=3. * indicated p<0.05 by Student’s t-test. Error bars show standard deviation.

**Supplemental Figure 9.**
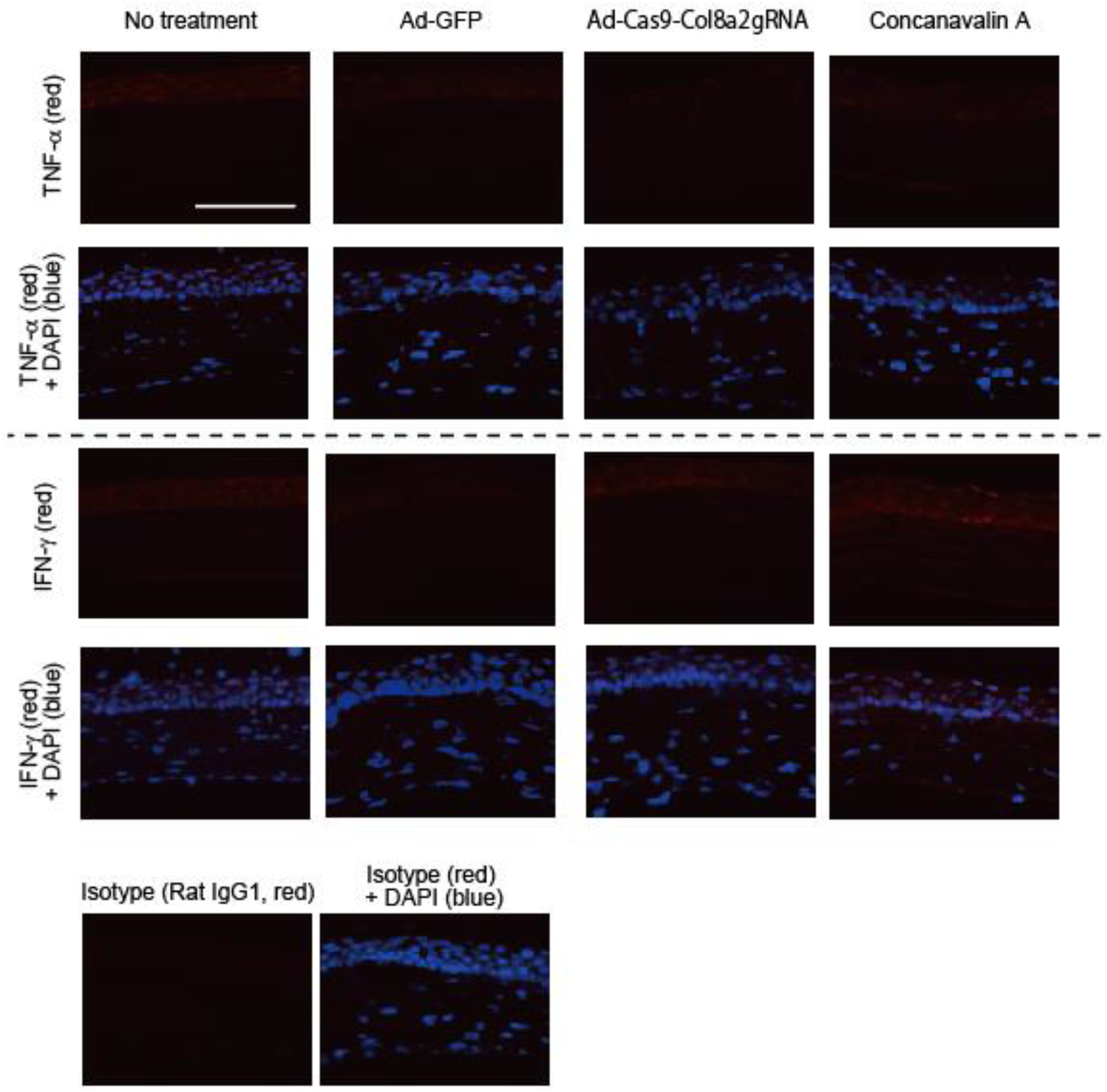
Intracameral injection of 0.25 × 10^8^ Ad-Cas9-Col8a2gRNA does not show significant induction of inflammation markers. TNFα and IFNγ were stained 4 weeks post Ad-GFP, Ad-Cas9-Col8a2gRNA or Concanavalin A (1μg) intravitreal injection. Scale bar is 100μm.

**Supplemental Figure 10.**
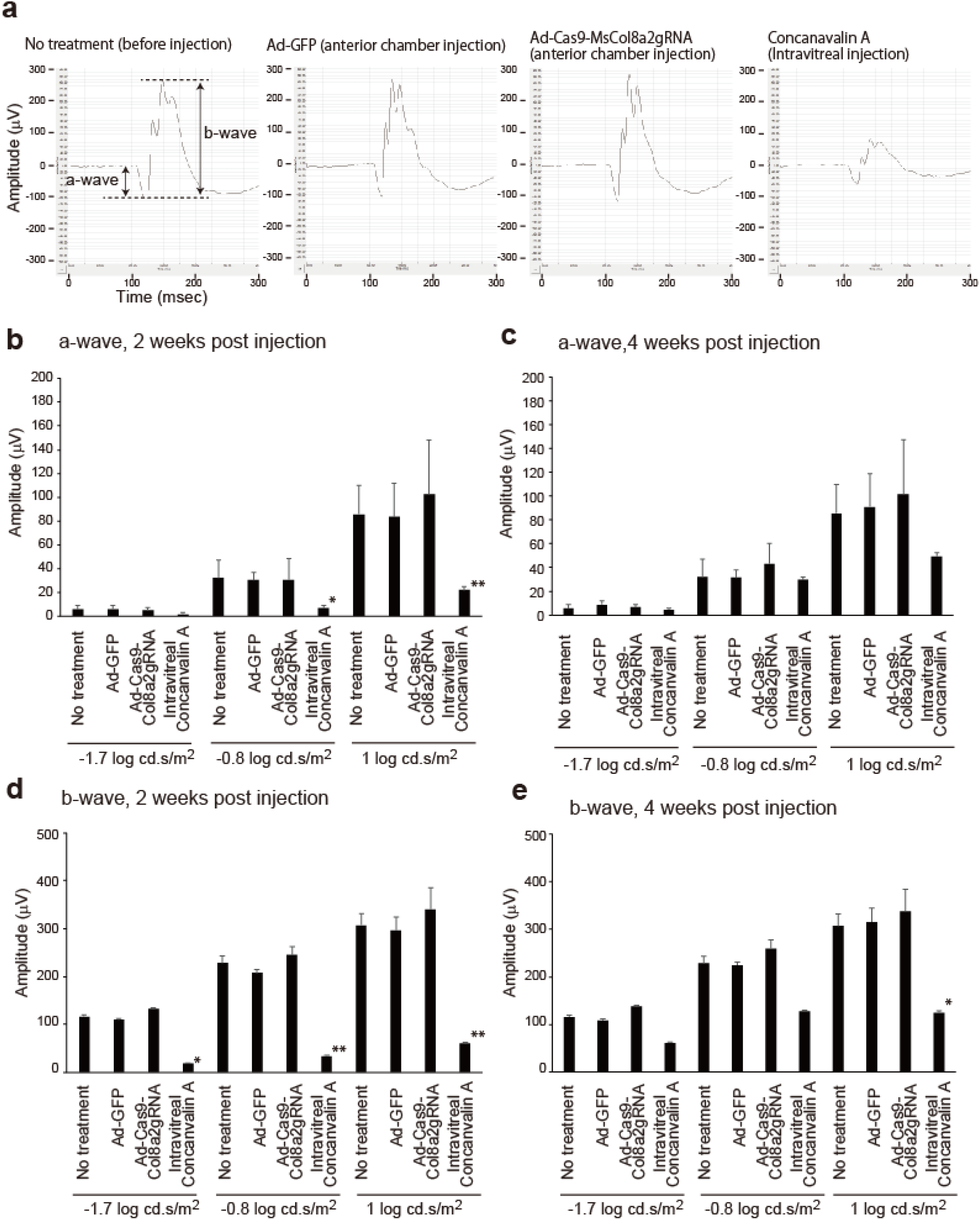
Ad-Cas9-Col8a2gRNA does not induce retinal disfunction. Dark adapted ERG was used for evaluation of retinal function. **a.** Representative ERG of no treatment (prior to injection), Ad-GFP (anterior chamber injection), Ad-Cas9-Col8a2gRNA (anterior chamber injection) and Concanavalin A (intravitreal injection). Intravitreal injection of Concanavlin A was used for postivie control by inducing retinal inflammation. **b, c**. a-wave of no treatment and each treatment 2 and 4 weeks post injection. We used three different stimulus light intensities (−1.7, −0.8 and 1 log cd.s/m^2^). **d, e.** b-wave of no treatment and each treatment 2 and 4 weeks post injection. n = 14 (no treatment), 6 (Ad-GFP), 6 (Ad-Cas9-Col8a2gRNA) and 2 (Concanavalin A, 1μg). * and ** indicated p<0.05 and 0.01 by student t-test compared to no treatment control.

**Supplemental Figure 11.**
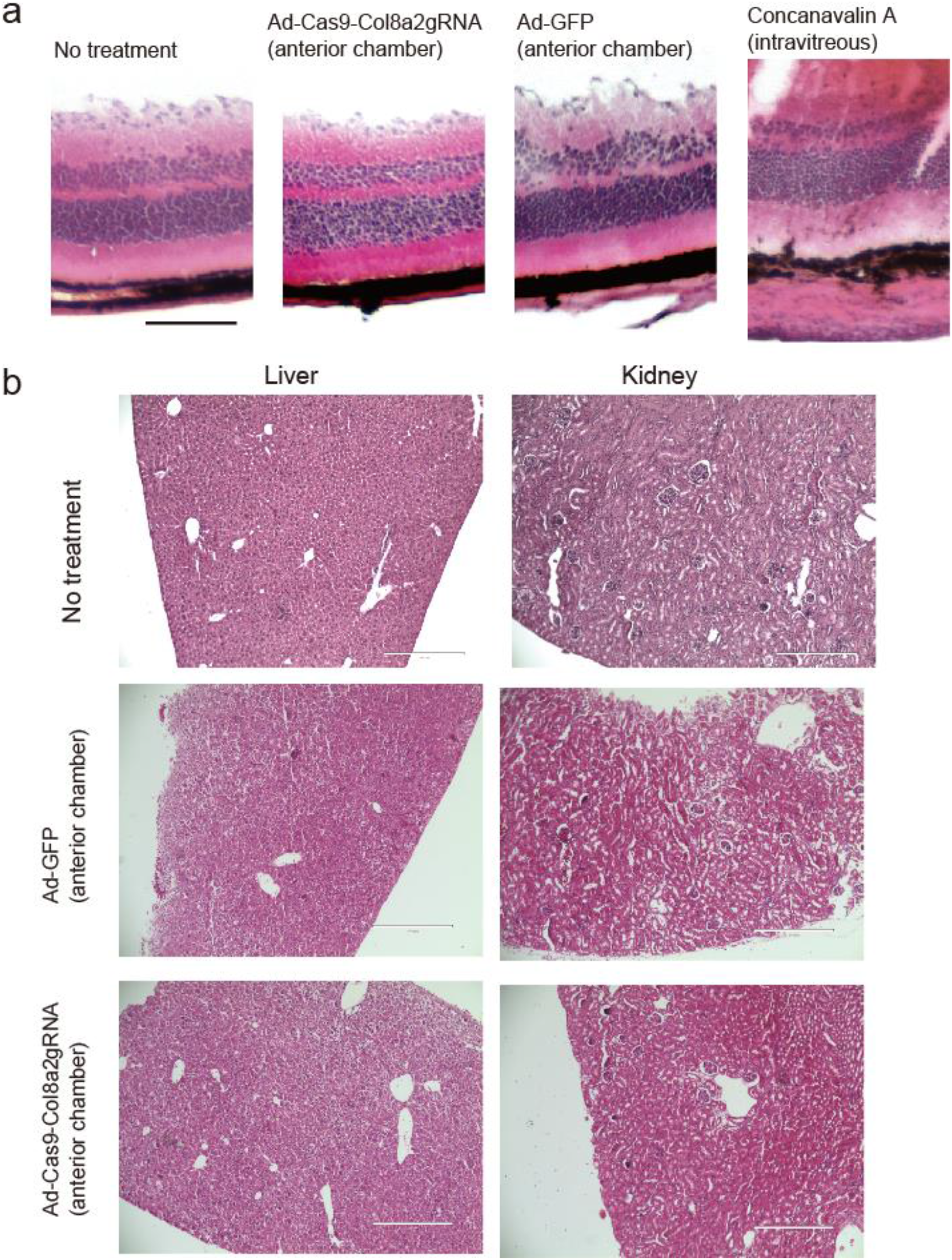
Anterior chamber injection of 0.25×10^8^ Ad-Cas9-Col8a2gRNA does not show retina, liver and kidney toxicity. 4 weeks post injection, we observed each tissue by HE staining. **a.** retina. Scale bar is 100μm. **b.** liver and kidney. Scale bar is 400μm.

**Supplemental Figure 12.**
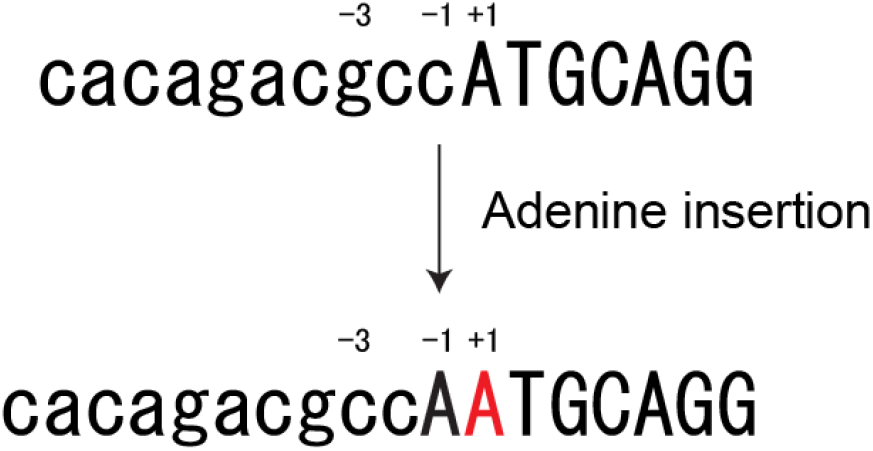
Single adenine insertion at the mouse *Col8a2* start codon. An adenine insertion produced a cryptic ATG codon that resulted in disruption of kozak sequence (g to c at -3 position).

**Supplemental Figure 13.**
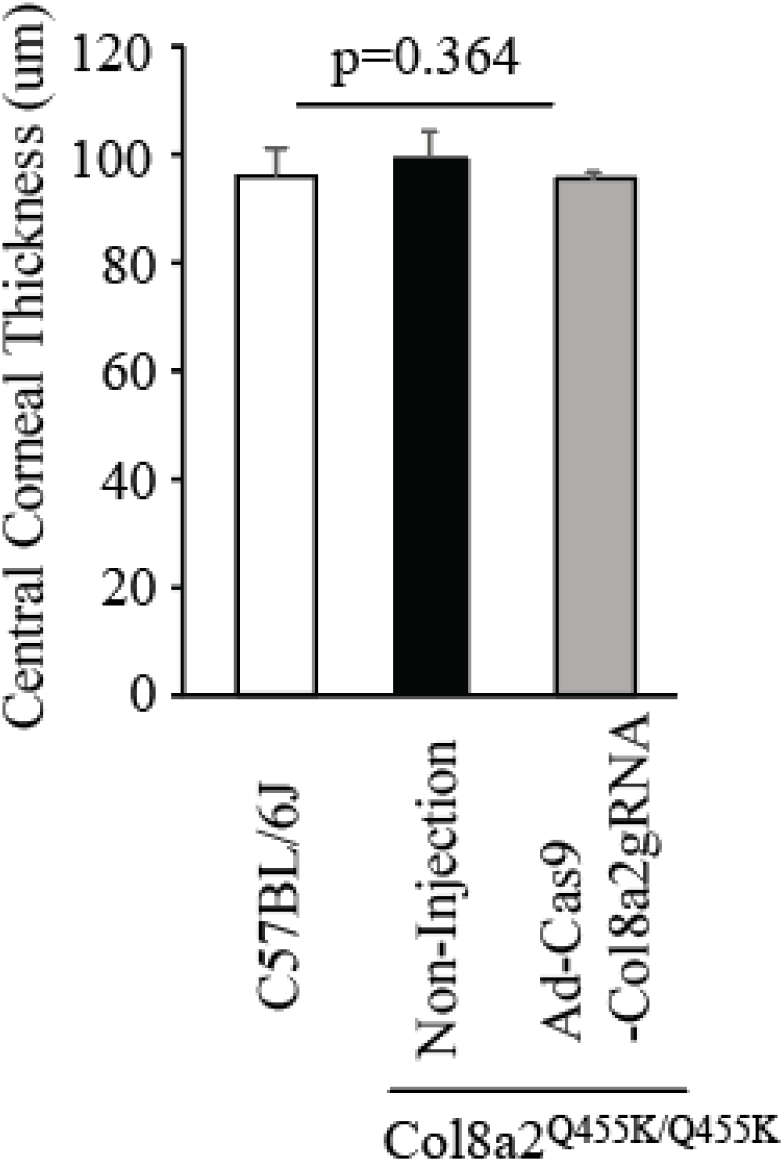
C*o*l8a2Q455K^/Q455K^ mice (12 months old) did not show a significant difference in central corneal thickness. Central corneal thickness was measured by corneal OCT, p-value was calculated by ANOVA.

**Supplemental Figure 14.**
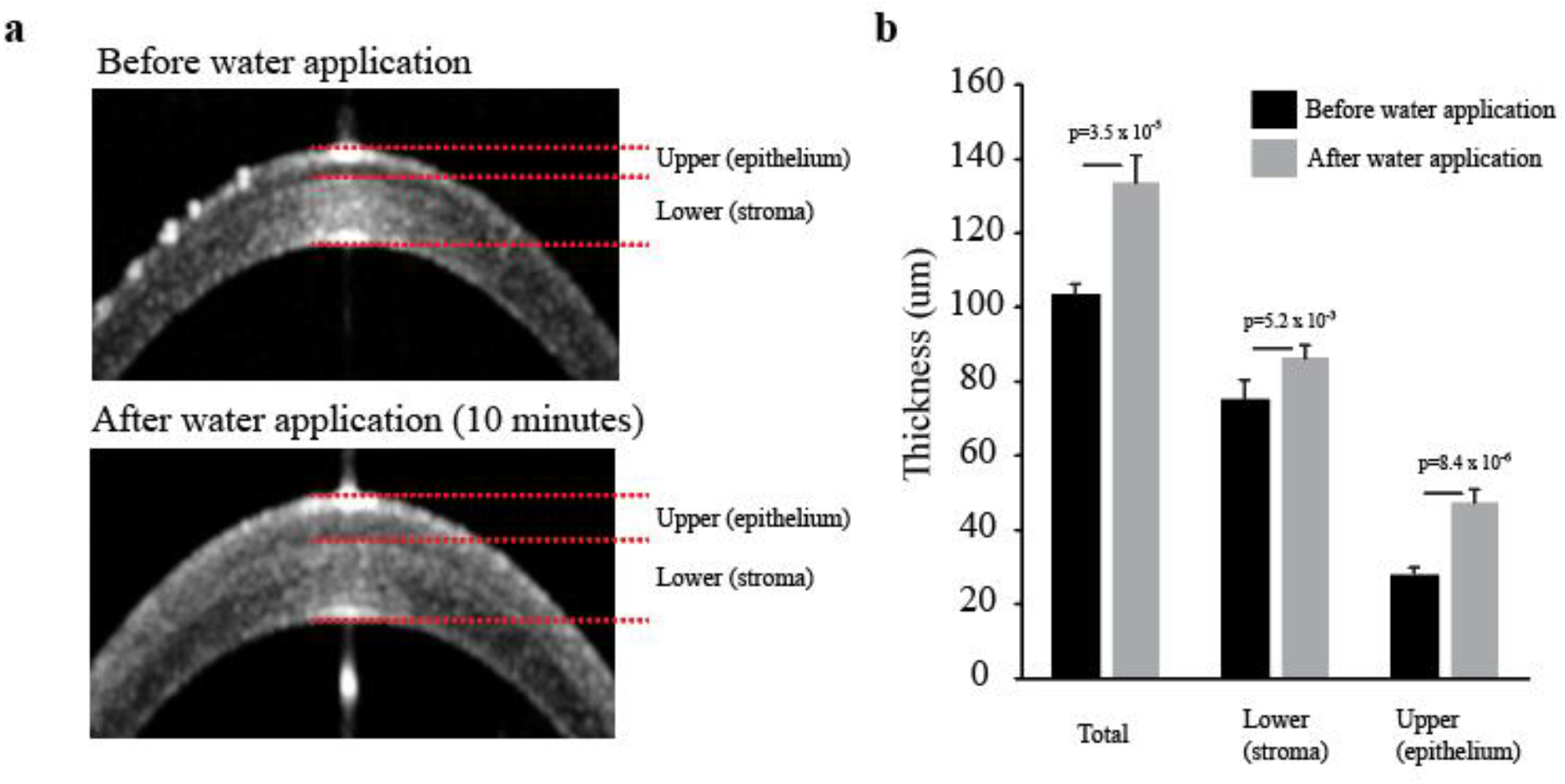
Water applied to the corneal surface expands the thickness of corneal epithelium rather than the stroma. **a,** Representative corneal OCT images before and after water was applied for 10 minutes. **b,** The average thickness of total, upper (epithelium) and lower (stroma) before and after water application for 10 minutes. n=5. Error bars show standard deviation, p-value was calculated by Student’s t-test.

**Supplemental Figure 15.**
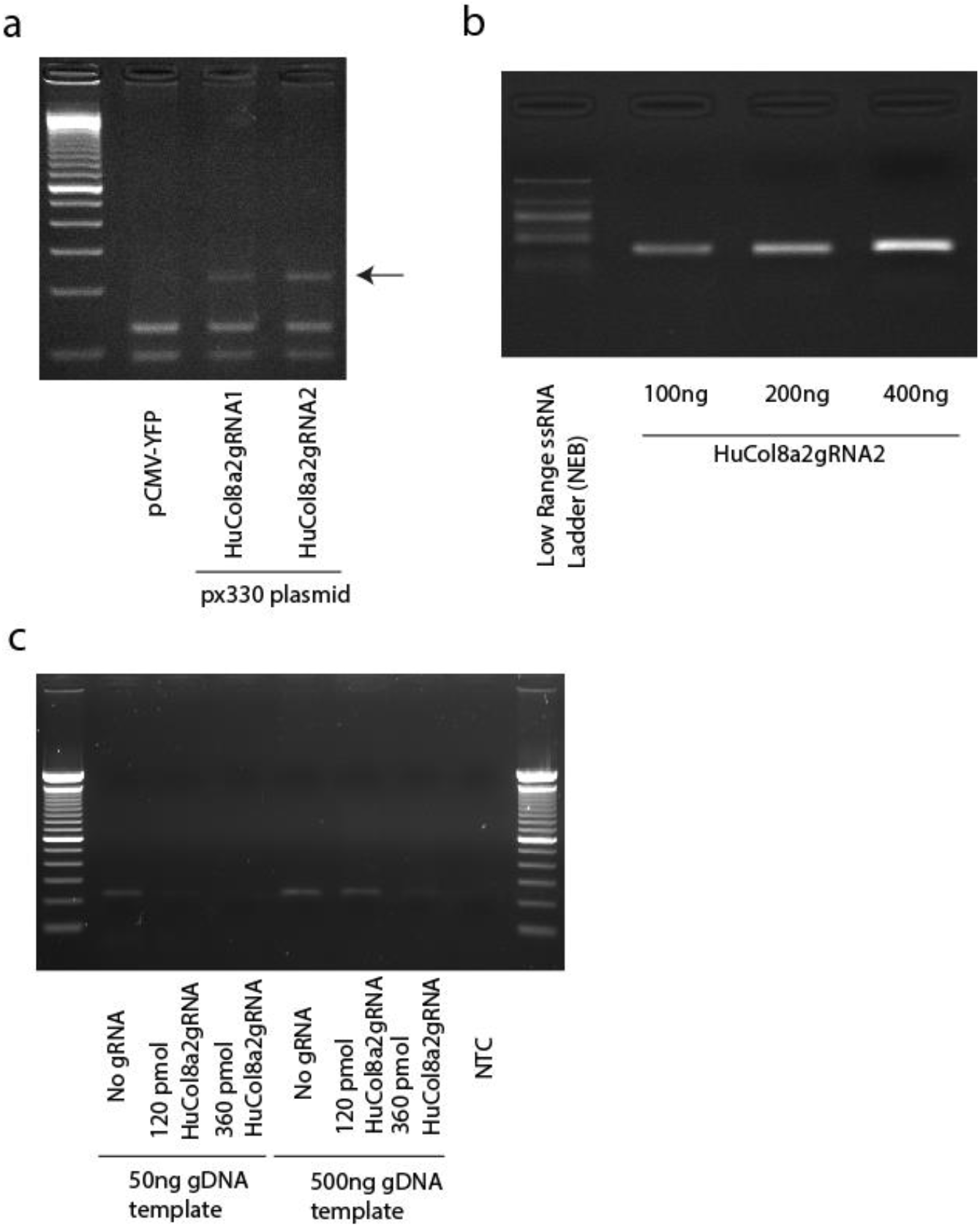
I*n vitro* digestion by Cas9/HuCol8a2gRNA. **a**. Plasmid based Cas9/HuCol8a2gRNA induced indels in AD293 cells. Arrow indicates indel band. **b.** Gel electrophoresis image of *in vitro* transcription of HuCol8a2gRNA. **c,** PCR confirmed *in vitro* digestion of purified AD293 genomic DNA by Cas9/HuCol8a2gRNA. PCR primers were designed to target the digestion site.

**Supplemental Table 1.**
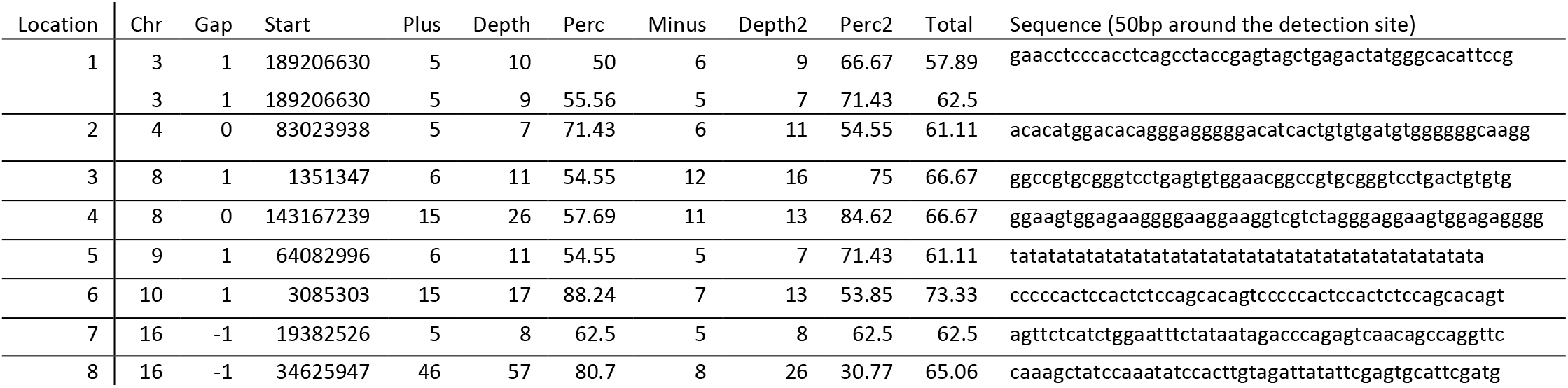
HuCol8a2gRNA off-target sites without homology.

**Supplemental Table 2.**
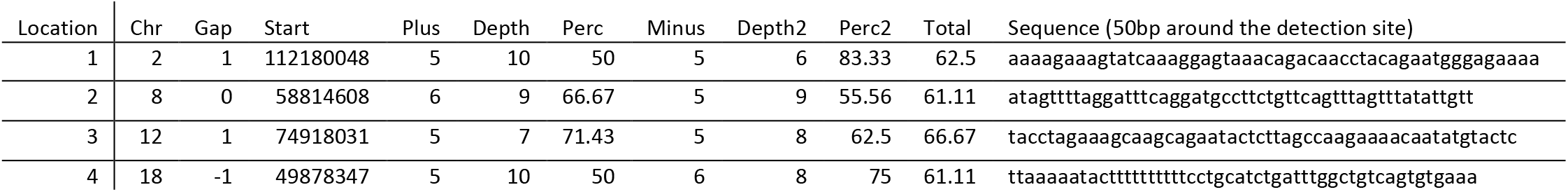
Detected sites with digenome score >60 in the control genomic DNA.

